# Microbiome responses to natural *Fusarium* infection in field-grown soybean plants

**DOI:** 10.1101/2025.03.27.645697

**Authors:** Sietske van Bentum, Bridget S. O’Banion, Alexandra D. Gates, Hans A. Van Pelt, Peter A.H.M. Bakker, Corné M.J. Pieterse, Sarah L. Lebeis, Roeland L. Berendsen

## Abstract

The rhizosphere microbiome influences plant health by mediating plant-pathogen interactions. Plants can recruit protective microbes in response to disease, but the consistency of this process in field conditions is unclear. We examined the rhizosphere microbiome of field-grown soybean (*Glycine max* L.) naturally infected with root pathogens across three commercial fields in Kentucky, USA. Symptomatic and asymptomatic plants were sampled to assess disease-associated shifts in the rhizosphere microbiome. Amplicon sequencing identified a diverse *Fusarium* community, with one *Fusarium solani* amplicon sequence variant (ASV) consistently enriched in diseased plants, identifying it as the likely pathogen. While microbial communities differed between diseased and healthy plants, these shifts were largely field-specific. Several fungal ASVs with known biocontrol potential (*Clonostachys rosea, Penicillium*, and *Trichoderma*) were enriched in healthy plants, implying a role in disease suppression. A *Sphingomonas* ASV, a genus previously linked to plant protection, was more abundant in diseased plant rhizospheres in two fields, suggesting pathogen-triggered recruitment. Conversely, *Macrophomina phaseolina*, a generalist root pathogen, was enriched in the rhizosphere of diseased plants in all fields, indicating possible co-infection with *F. solani*. These findings reveal complex pathogen-microbe interactions in field conditions and emphasize the need for field-specific microbiome research to inform sustainable disease management strategies.

## Introduction

Crop pests and pathogens are commonly controlled with agrochemicals that can persist in and damage the environment (Mahmood *et al*., 2016; Meena *et al*., 2020; Woodcock *et al*., 2017). Moreover, agrochemicals are mostly ineffective against soilborne pathogens. While plant-beneficial microbes that protect plants against attackers are considered more sustainable alternatives to chemical protection, their efficacy is often inconsistent across different environments (Lutz *et al*., 2023; Owen *et al*., 2015; Poppeliers *et al*., 2023; Thilakarathna & Raizada, 2017). Plants can play a key role in improving the effectiveness of plant-beneficial microbes by recruiting them from the soil environment in response to pathogen infection (Berendsen *et al*., 2018; Liu *et al*., 2021; Yin *et al*., 2021; Yuan *et al*., 2018). For instance, plant resistance-inducing bacteria were found to increase in abundance in the rhizosphere of the model plant *Arabidopsis thaliana* following infection by the foliar pathogen *Hyaloperonospora arabidopsidis* (Berendsen *et al*., 2018; Goossens *et al*., 2023). The enrichment of these protective rhizobacteria appears to involve a disease-induced change in the root exudation profile, suggesting that plants actively promote the beneficial microbes that come to their rescue (Goossens *et al*., 2023; Vismans *et al*., 2022; Yuan *et al*., 2018). Moreover, plant-protective bacteria can persist in soil and suppress disease in subsequent populations of plants grown in the same soil (Bakker *et al*., 2018; Berendsen *et al*., 2018; Goossens *et al*., 2023; Luo *et al*., 2022; Vismans *et al*., 2022; Yin *et al*., 2021; Yuan *et al*., 2018).

Disease-associated enrichment of plant-beneficial rhizosphere microbes has been described for several crop species, such as wheat, potato and sugar beet (Carrión *et al*., 2019; Liu *et al*., 2021; Mendes *et al*., 2011; Yin *et al*., 2021), and it is thought that disease suppressive soils arise from these processes (Spooren *et al*. 2024). Currently, it is unknown whether leguminous plants also recruit beneficial microbes in response to pathogen infection, although soil microbes can suppress the incidence and severity of *Fusarium*-associated sudden death syndrome (SDS) in soybean (Westphal & Xing, 2011).

Soil microbial communities in SDS-affected field patches host phylogenetically diverse bacteria and fungi, including potential plant-beneficial taxa (Srour *et al*., 2017). Similarly, natural infections by *Septoria glycines* or *Phytophthora sojae* in field-grown soybean plants altered rhizosphere bacterial and fungal communities (Díaz-Cruz & Cassone, 2022). Sampling of plants across the growing season revealed differentially abundant fungal taxa between rhizospheres of infected and healthy plants, although specific microbial patterns varied by disease and growth stage. This highlights that spatial and temporal variation in the microbiome of field-grown plants can complicate the identification of candidate beneficial or detrimental taxa in the rhizosphere.

To examine the consistency of disease-associated microbiome dynamics and potential recruitment of beneficial microbes in soybean, we studied the rhizosphere microbiome of naturally-infected soybean plants affected by a soilborne *Fusarium* species in three commercial fields in Kentucky (USA). Using DNA amplicon sequencing, we compared the bacterial and fungal rhizosphere communities of healthy and diseased plants. Recognizing that different fields differ in the composition of their resident soil microbiomes (Tkacz *et al*., 2020; Walters *et al*., 2018), we sampled plants across multiple soybean fields to identify specific microbial taxa consistently associated with plant health or disease, independent of local environmental variation. Our goal was to uncover soybean-adapted candidate beneficial microbes enriched during pathogen infection across diverse environments, offering potential to support soybean resilience against disease.

## Materials and Methods

### Sampling procedures of field-grown soybean plants

In each soybean field, thirty healthy and thirty plants with leaf chlorosis, leaf necrosis and defoliation were sampled. Of note, petioles of fallen leaves remained attached to the plant. The disease occurred in patches, thus symptomatic plants were sampled from these patches. Each healthy plant was sampled at variable distance from any sampled symptomatic plant. This was recorded along with the GPS coordinates for each sampled plant (fig. S1). Foliar symptoms were scored for each plant and a trifoliate leaf with the most severe symptoms based on five classes (0-4). For whole plant scoring: 0 = no symptoms; 1 = <10% of trifoliate leaves showed symptoms; 2 = 10-25% of trifoliate leaves showed symptoms; 3 = 25-50% of trifoliate leaves showed symptoms; 4 = >50% of trifoliate leaves showed symptoms. For individual trifoliate leaves: 0 = no symptoms; 1 = a single small lesion or spot of chlorosis that covers up to 1% of leaf surface; 2 = 2-9% of leaf surface covered in lesions and/or chlorosis; 3 = 10-25% of leaf surface covered in lesions and/or chlorosis; 4 = >25% of leaf surface covered in lesions and/or chlorosis. All subsequent steps were performed while wearing ethanol-cleaned gloves. The trifoliate leaf used for scoring disease severity was placed inside a ziplock bag (Whirl-Pak, Pleasant Prairie, United States) and stored on ice until transferal to a 50-ml Falcon tube (Greiner, Kremsmünster, Austria) and stored at -20°C later the same day. Each plant was uprooted with a shovel to 20-25 cm depth. Roots were cleaned of excess soil and stored in ziplock bags (Whirl-Pak, Pleasant Prairie, United States) on ice until processing later the same day. Bulk soil was sampled ten times with a soil corer inside each field and packed in 50-ml Falcon tubes that were stored on ice until storage later the same day. When roots were processed, lateral roots were cut with ethanol-cleaned scissors and transferred to 50-ml Falcon tubes. All tubes with leaf, root and soil material were stored long-term at -20°C before and after transportation on dry ice to Utrecht (the Netherlands).

### DNA extractions on bulk soil and soybean root samples

DNA was extracted in Utrecht (NL) from root and soil samples based on an adapted protocol of the DNeasy PowerSoil kit (Qiagen, Hilden, Germany). This kit is designed for small biological samples (up to 0.25 g) and was adjusted to enable extractions from soybean root samples (approx. 0.1 – 12 g per sample). A detailed protocol is available in the supplemental methods. Bulk soil samples were subsampled by taking approx. 0.25 g per sample to a sterile 2-ml Eppendorf tube (Eppendorf, Hamburg, Germany). Two sterile glass beads (⍰ mm) were added to each subsampled soil sample and ten beads to each root sample. Soil and root samples were incubated at 70°C for 10 min prior to cell lysis. Soil samples were physically disrupted in a TissueLyser II (Qiagen, Hilden, Germany) in a mixture of 0.75 ml bead solution and 0.06 ml C1 solution. The TissueLyser II was run twice for 10 min at 30 Hz. Root samples were disrupted in a SK550 1.1 heavy-duty paint shaker (Fast & Fluid, Sassenheim, the Netherlands) in a similar mixture of bead solution and C1 solution (3 : 0.24 ratio). The total volume of lysis solution was adjusted to root sample weight to obtain sufficient supernatant without soil particles, with a minimal total volume of 11 ml. The paint shaker was set at 270 sec at speed 3 followed by 270 sec at speed 6. In this setup, the roots remained largely intact, suggesting that DNA was primarily extracted from the outside of the roots (the rhizosphere) rather than from the root endosphere. After cell lysis, 700 μl supernatant per sample was transferred to a new 2-ml Eppendorf tube. Supernatant from soil and root samples was cleaned with C2 and C3 solutions as described in the protocol of the DNeasy PowerSoil kit. DNA was subsequently purified based on the protocol of the MagMAX Microbiome Ultra Nucleic Acid Isolation kit (ThermoFisher, Waltham, United States) with a KingFisher Flex Purification System (ThermoFisher, Waltham, United States). Specifically, 500 μl supernatant per sample was combined with 500 μl binding bead mix in a 96-well plate, the latter mix consisting of binding solution and ClearMag beads (45 : 2 ratio). The DNA was subsequently washed in 96-well plates in the KingFisher Flex Purification System, twice in washing buffer (Tris 7.5 mM, NaCl 97.5 mM, ethanol 50%, Milli-Q) and twice in 80% ethanol. After washing, beads with DNA were air-dried in the machine for 8 min before DNA was eluted in 50 μl Tris (100 mM in MQ, pH 8.0-8.5). The concentration and quality of DNA was measured with Nanodrop 2000/2000c (ThermoFisher, Waltham, United States) and Qubit Fluorometer 3.0 with a Qubit dsDNA BR Assay kit (Invitrogen, Waltham, United States).

### 16S rRNA gene and ITS2 amplicon library preparations and sequencing

DNA extracted from bulk soil and soybean rhizosphere samples was sent to Génome Québec (Québec, Canada) for 16S and ITS2 amplicon library preparations and sequencing. This totaled 10 soil, 30 diseased plant and 29 healthy plant samples in Field 1; 10 soil, 30 diseased plant and 30 healthy plant samples in Field 2; and 10 soil, 28 diseased plant and 28 healthy plant samples in Field 3. Amplicons were sequenced as paired-end 250bp sequences on an Illumina NovaSeq 6000 SP. Blocking primers were used to prevent the amplification of plant-derived DNA (Agler *et al*., 2016; Lundberg *et al*., 2013). The sequences of amplicon and blocking primers can be found in table S1.

### Amplicon sequencing data processing

The 16S and ITS2 amplicon sequences from all three fields combined were initially processed in R (v4.2.2; R Core Team, 2022). The sequences had been demultiplexed at the sequencing facility. Amplicon primer sequences were removed with cutadapt (Martin, 2011). Forward and reverse reads were filtered and ASVs were determined with DADA2 (Callahan *et al*., 2016). ITS2 amplicon reads were filtered with maximum 2 expected errors and minimum read length = 50 nt. 16S amplicon reads were trimmed based on their quality score (Q > 30), filtered with maximum 2 errors and truncated at 225 or 220 nt for forward and reverse reads, respectively. Reads that matched against the phiX genome were removed. Due to the binned quality scores obtained from the Illumina NovaSeq platform, sequencing errors were estimated based on a modified loess fit function where weights, span and degree were altered and monotonicity was enforced (Oliverio & Holland-Moritz, 2021). Forward and reverse reads were merged and chimeras removed. Contaminant reads were removed based on occurrence in true samples and blanks with the decontam package (Davis *et al*., 2018). Taxonomy was assigned to ASVs with BLAST+ local alignment in Qiime2 (version 2022.11; Bolyen *et al*., 2019). Fungal taxonomy was assigned to ITS2 ASVs based on the UNITE 8.3 database (Nilsson *et al*., 2019). Non-fungal ASVs were removed, including plant, Rhizaria, Metazoa, Alveolata, Protista and unassigned reads. Bacterial taxonomy was assigned to 16S ASVs based on the SILVA 132 database (Quast *et al*., 2013). Non-bacterial ASVs were removed from the 16S datasets, including plant, Archaea and unassigned reads.

Based on cumulative abundances, rare ASVs were excluded if they were represented by <53 reads in the ITS2 dataset and by <195 reads in the 16S dataset. ASVs were also excluded if they were present in <4 samples in the fungal dataset or <27 samples in the 16S dataset. In the ITS2 dataset, samples with <1,061 reads were filtered to avoid an effect of low sequencing depth, excluding 9 samples from Field 1, 7 samples from Field 2, and 18 samples from Field 3. The final datasets comprised 187 fungal ASVs across 60 samples and 6,189 bacterial ASVs across 69 samples in Field 1; 219 fungal ASVs across 63 samples and 6,016 bacterial ASVs across 70 samples in Field 2; and 169 fungal ASVs across 48 samples and 6,269 bacterial ASVs across 66 samples in Field 3.

### Data analysis and visualization

The raw sequencing data have been deposited in the European Nucleotide Archive (ENA) at EMBL-EBI under accession number PRJEB87115 (https://www.ebi.ac.uk/ena/browser/view/PRJEB87115). Code and files from data analysis are provided at Github (https://github.com/SietskevB/FusariumUSA). Plots and analyses were mainly performed in R (v3.6.1; R Core Team, 2019). Boxplots, violin plots, histograms and stacked bar charts were created with ggplot2 (Wickham, 2016). The line plots showing the number of ASVs and sequencing depth per sample were created with vegan (Oksanen *et al*., 2020) and ggplot2 (Wickham, 2016). The heatmaps were created with tidyheatmap (Engler, 2022). The Fisher’s exact test was performed with package rstatix (Kassambara, 2021) and the Wilcoxon rank sum test with package stats (R Core Team, 2019). ITS2 sequences were extracted from NCBI GenBank and trimmed to match the region sequenced from the field samples (table S2; Clark *et al*., 2016). The ITS2 sequences from the field and GenBank were aligned in Qiime2 (version 2022.11) based on MAFFT and the rooted tree was visualized in iTOL (Bolyen *et al*., 2019; Letunic & Bork, 2021). Percentage identity scores were calculated with the MAFFT sequence analysis tool of EMBL-EBI (Madeira *et al*., 2022).

The ITS2 and 16S amplicon datasets were analyzed with phyloseq (McMurdie & Holmes, 2013) based on non-rarefied, relative read counts. Bray Curtis and Jaccard distances were calculated with phyloseq (McMurdie & Holmes, 2013). Differences in between-group distances were assessed with PERMANOVA in package pairwiseAdonis (Arbizu, 2017). The ordinations were plotted as Principal Coordinate Analysis (PCoA) with phyloseq (McMurdie & Holmes, 2013) and ggplot2 (Wickham, 2016).

Differentially abundant ASVs between healthy and diseased plants were determined based on five statistical tests: ANCOM-bc (Lin & Peddada, 2020), DESeq2 (Love *et al*., 2014), Fisher’s exact test, Simper analysis (Clarke, 1993) and Spearman correlations. ANCOM-bc was performed based on Lin (2020) using R packages microbiome (Lahti & Shetty, 2019) and nloptr (Johnson, 2020). DESeq2 was performed with package DESeq2 (Love *et al*., 2014). A pseudocount of 1 was added to the ITS2 amplicon data to execute the DESeq2 analysis. The Fisher’s exact test was performed with stats (R Core Team, 2019). Simper analysis was performed with stats and vegan (Oksanen *et al*., 2020; R Core Team, 2019). Spearman rank correlations were performed with jmuOutlier (Garren, 2019). ASVs were considered differentially abundant if the FDR-adjusted *p*□≤□0.05. Sparse ASVs denoted as structural zeroes by ANCOM-bc were ignored in downstream analysis unless also detected by at least one other statistical method. The relative abundance of each differentially abundant ASV was normalized by its average relative abundance across healthy plant rhizosphere samples and subsequently log-transformed with a pseudocount of 1. These log-transformed, healthy plant-normalized relative abundances of differentially abundant ASVs were plotted in heatmaps.

## Results

### Disease severity in three commercial soybean fields in Kentucky

To identify disease-associated differences in the rhizosphere microbiome of naturally-infected soybean plants, three commercial soybean fields in Kentucky (USA) were sampled. Symptomatic plants showed leaf chlorosis, leaf necrosis, and premature defoliation with petioles that remained attached to the plant. These symptoms match those caused by several *Fusarium* spp.: the root rot pathogen species *Fusarium solani* (Nelson, 2015) as well as four species that cause SDS: *Fusarium brasiliense, Fusarium crassistipitatum, Fusarium tucumaniae* and *Fusarium virguliforme* (Aoki *et al*., 2005; Hartman *et al*., 2015b; Wang *et al*., 2019). Disease severity was scored as the percentage of trifoliate leaves with symptoms of infection per sampled plant (fig. 1A) and as the extent of foliar symptoms in the most strongly affected trifoliate leaf per plant (fig. 1B). Because disease symptoms were absent in healthy plants, they were not scored. In each field, all degrees of disease severity at the leaf and whole-plant level were observed in the sampled symptomatic plants. The disease severity of symptomatic plants was similar between all three fields at the plant and single leaf level (Fisher’s exact test, adjusted *p* > 0.05).

**Figure 1.**
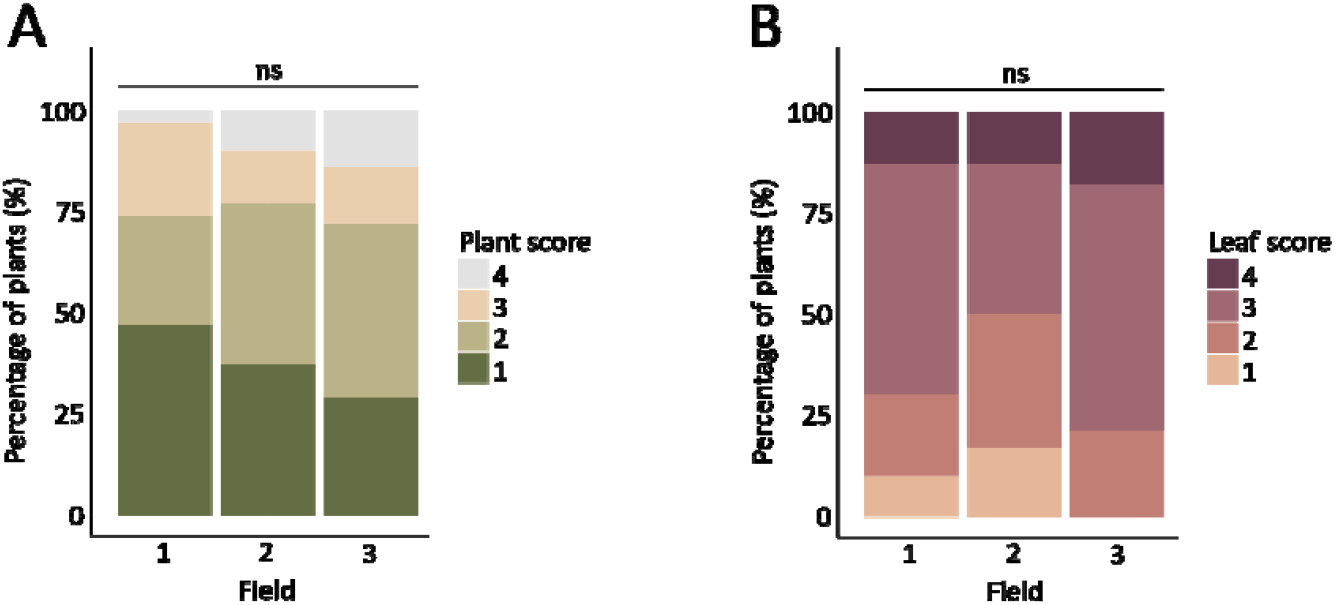
Disease severity of symptomatic plants sampled in three commercial soybean fields. **A)** Disease severity in plants with symptoms of disease: 1 = <10% of trifoliate leaves showed symptoms; 2 = 10-25% of trifoliate leaves showed symptoms; 3 = 25-50% of trifoliate leaves showed symptoms; 4 = >50% of trifoliate leaves showed symptoms. **B)** Disease severity in the most strongly affected trifoliate leaf of each plant with symptoms of disease: 1 = a single small lesion or spot of chlorosis that covers up to 1% of leaf surface; 2 = 2-9% of leaf surface covered in lesions and/or chlorosis; 3 = 10-25% of leaf surface covered in lesions and/or chlorosis; 4 = >25% of leaf surface covered in lesions and/or chlorosis. Data based on 30 diseased plants in Field 1; 30 diseased plants in Field 2; and 28 diseased plants in Field 3. No significant differences were observed based on Fisher’s exact test (adjusted *p* > 0.05).

### Microbial diversity in the rhizosphere of field-grown soybean plants

To investigate the impact of disease on the soybean rhizosphere microbiome, bacterial and fungal communities associated with bulk soil and roots from healthy and diseased plants were characterized based on 16S rRNA gene and ITS2 amplicon sequencing, respectively. The fungal ITS2 dataset comprised 187 amplicon sequence variants (ASVs) across 60 samples in Field 1; 219 ASVs across 63 samples in Field 2; and 169 ASVs across 48 samples in Field 3. The bacterial 16S dataset comprised 6,189 ASVs in Field 1; 6,016 ASVs in Field 2; and 6,269 ASVs in Field 3. The majority of ITS2 and 16S ASVs was detected in all three fields, and these shared ASVs comprised 103 fungal ASVs and 5,724 bacterial ASVs.

Across the three fields, the average sequencing depth for 16S and ITS2 amplicons did not differ between healthy and diseased plants (fig. S2; Wilcoxon signed rank test, *p* > 0.05). Moreover, the sequencing depth was sufficient to capture the biological diversity in all samples, since the number of ASVs detected in each sample reached saturation (fig. S3). The number of fungal ASVs detected per sample ranged from 5 to 39, which suggests a relatively low fungal diversity in each sample. This contrasts the bacterial data where 425 to 2,432 ASVs were detected per sample. Since the bacterial and fungal data were obtained from the same field samples, the difference in the total number of bacterial and fungal ASVs in each field likely reflects biological rather than technical variation.

The average number of bacterial ASVs did not differ between healthy and diseased plant rhizosphere samples in any of the three fields (fig. S4A; Wilcoxon signed rank test, *p* > 0.05). The average number of fungal ASVs was significantly higher in the rhizosphere of diseased plants in fields 2 and 3, but not in Field 1 (fig. S4B; Wilcoxon signed rank test, *p* < 0.05). Although the number of ASVs was low and variable across samples, the slight increase in number of fungal ASVs in the rhizosphere of diseased plants in fields 2 and 3 suggests potential shifts in microbiome composition associated with plant disease.

Although 103 fungal ASVs were detected across all three fields, the majority of fungal ASVs were detected in only one or two samples per field (fig. S5). Sparsity of microbial features is common in microbiome datasets and needs to be carefully considered in downstream analyses (Nearing *et al*., 2022).

### Disease has a minor effect on soybean rhizosphere microbiome composition

To determine the effect of disease on the soybean rhizosphere microbiome, we performed principal coordinate analysis (PCoA). The composition of the bacterial communities was clearly affected by field, as confirmed by permutational analysis of variance (PERMANOVA; R^2^ > 0.03, adjusted *p* < 0.01; fig. S6A, table S3). The bacterial communities of Field 2 were especially dissimilar from Field 1 and 3. The field effect was smaller for fungal community composition yet still significant (PERMANOVA, R^2^ = 0.02, adjusted *p* < 0.01; fig. S6B, table S4). Rhizosphere bacterial communities were clearly dissimilar from bulk soil communities in all three fields (PERMANOVA; R^2^ > 0.15, adjusted *p* < 0.01) while this distinction was much less pronounced in the fungal communities (PERMANOVA; R^2^ = 0.01-0.02, adjusted *p* ≤ 0.58; fig. S6CD, table S3-S4). This suggests that the soybean rhizosphere environment was more selective for soil bacteria than for soil fungi.

Disease incidence did not significantly (*p <* 0.05) impact rhizosphere bacterial communities across all fields combined, although a trend was observed (PERMANOVA, adjusted *p* = 0.06; fig. S6C, table S3). At the individual field level, bacterial rhizosphere communities of healthy and diseased plants differed significantly in composition only in Field 1 (PERMANOVA, R^2^ = 0.04, adjusted *p* = 0.01; fig. 2ABC, table S5). Fungal rhizosphere communities of diseased plants were slightly but significantly different from healthy plants across the three fields combined (PERMANOVA, R^2^ = 0.01, adjusted *p* = 0.03; fig. S6D, table S4). However, at the field level, fungal communities were only significantly different in Field 3 (PERMANOVA, R^2^ = 0.040, adjusted *p* < 0.01; fig. 2DEF, table S6).

**Figure 2.**
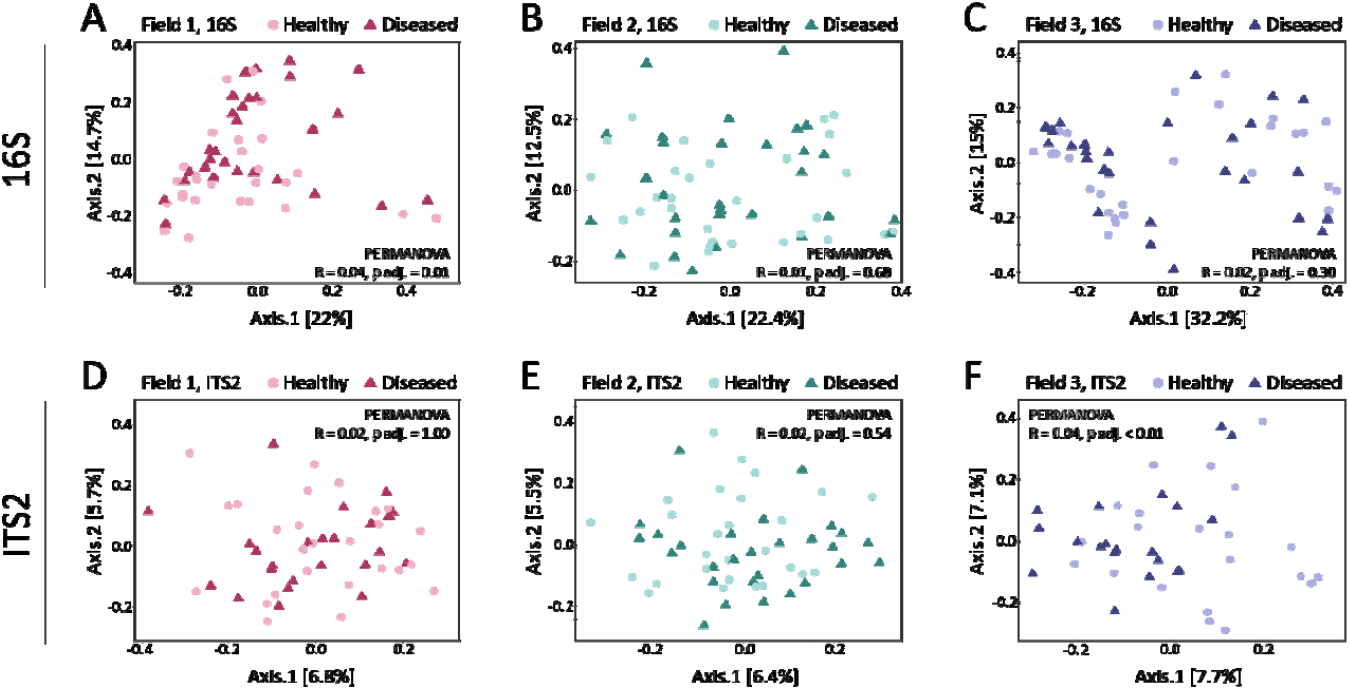
Principal coordinate analysis (PCoA) plots of bacterial and fungal communities of soybean rhizosphere samples from three fields. **(A–C)** PCoA plots based on Bray-Curtis distances showing bacterial rhizosphere communities for Field 1 (**A**), Field 2 (**B**), and Field 3 (**C**). **(D-F)** PCoA plots based on Jaccard distances showing fungal rhizosphere communities for Field 1 (**D**), Field 2 (**E**), and Field 3 (**F**). Light circles represent the rhizosphere samples of healthy plants and dark triangles those of diseased plants. Statistical analyses of the differences between microbial communities are provided in tables S1 and S2.

In conclusion, disease had a minor effect on the composition of the soybean rhizosphere microbiome, with significant differences on bacterial rhizosphere communities in Field 1 and fungal rhizosphere communities in Field 3.

### Differentially abundant microbial ASVs between the rhizosphere of healthy and diseased plants

To identify bacterial and fungal ASVs affected by disease in field-grown soybean plants, differentially abundant ASVs between healthy and diseased plants were identified using an approach similar to Vismans *et al*. (2022). Because different statistical tests detect varying numbers and identities of differentially abundant ASVs (Nearing *et al*., 2022), we applied five complementary statistical methods to capture a diverse set of ASVs: ANCOM-bc, DESeq2, Fisher’s exact test, Simper analysis and Spearman rank correlations (Lin & Peddada, 2020; Love *et al*., 2014; Clarke, 1993).

This approach identified 43 differentially abundant bacterial ASVs in field 1, the field with a significant effect of disease on rhizosphere bacterial communities (fig. 2A & 3). Among these, five *Enterobacter* ASVs were significantly enriched in the rhizosphere of diseased plants. Notably, *Enterobacter cloacae* has been previously implied in *in vitro* antifungal activity against *F. oxysporum* and in enhanced plant resistance against *Fusarium* wilt in spinach and maize (Ravi *et al*., 2022; Sallam *et al*., 2024; Tsuda *et al*., 2001). A *Sphingomongas* ASV 37efc was also enriched on diseased plants in Field 1. In contrast, three *Pelomonas* ASVs and five *Burkholderiaceae* ASVs were more abundant in the rhizosphere of healthy plants in this field. There were no differentially abundant bacterial ASVs in Field 2 and eight differentially abundant bacterial ASVs in Field 3 (fig. S7). The differentially abundant ASVs in Field 3 included the same *Sphingomonas* ASV 37efc that was differentially abundant in Field 1 (fig. 3, fig. S7). This bacterial genus has been previously connected to disease suppressiveness against black root rot of tobacco caused by *Thielaviopsis basicola* and bacterial wilt of tomato plants caused by *Ralstonia solanacearum*, however, a link to plant-pathogenic *Fusarium* species has not yet been established (Kyselková *et al*., 2014; Wei *et al*., 2019). *Sphingomonas* 37efc was also present at low abundance but not differentially abundant between healthy and diseased plants in Field 2. The other seven differentially abundant ASVs in Field 3 were phylogenetically diverse and were not differentially abundant in Field 1.

**Figure 3.**
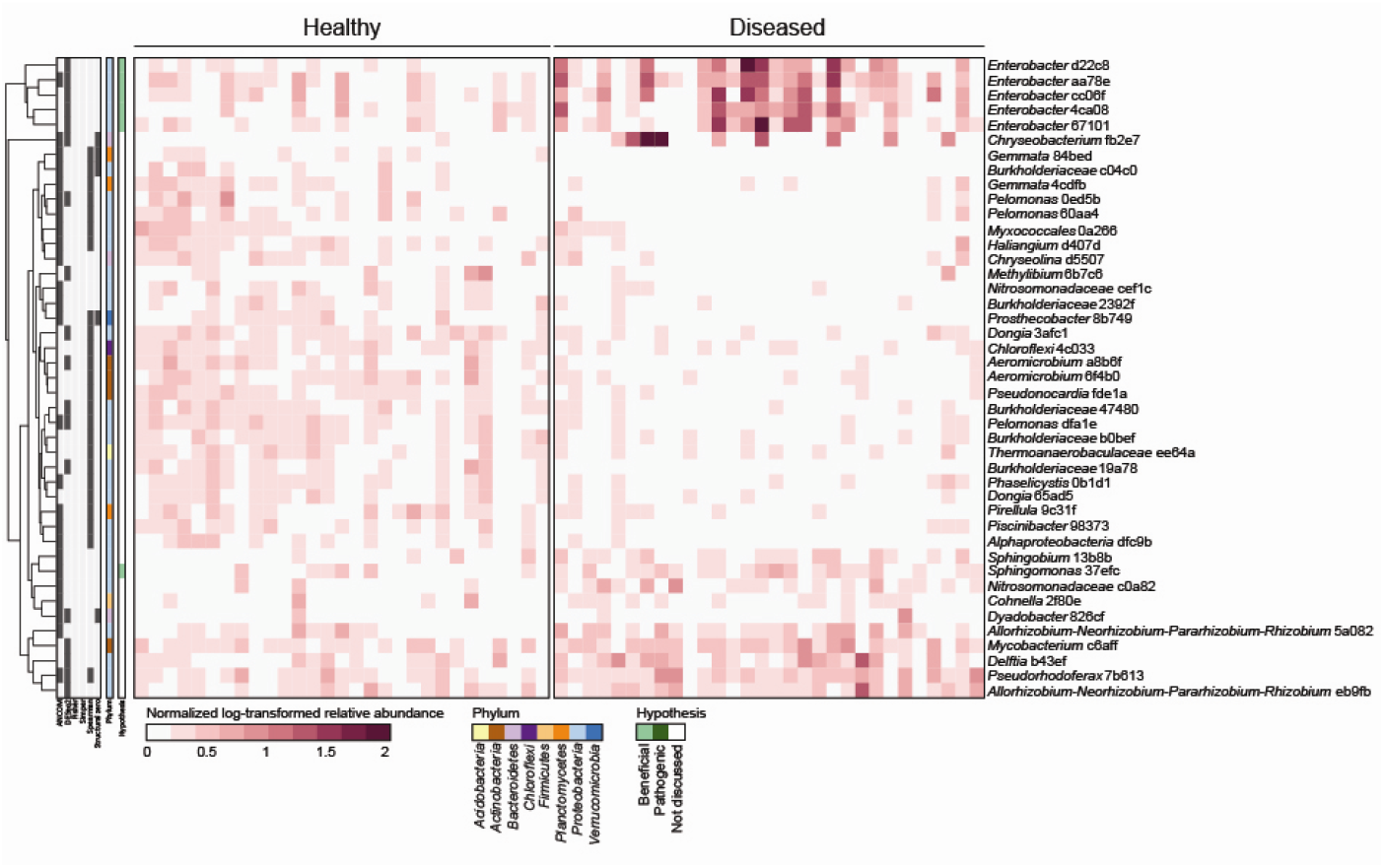
Forty-three differentially abundant bacterial ASVs in the rhizosphere of healthy and naturally-infected soybean plants in Field 1. The heatmap presents log-transformed relative abundances normalized by the average relative abundance of each ASV across all healthy plant samples. Colors to the left of the heatmap indicate whether an ASV or its associated taxon is discussed in the text (green), the phylum of each ASV (multicolor), and the statistical method(s) by which the ASV found differentially abundant (grey). Clustering of ASVs is based on Euclidean distances.

While we also applied the five statistical tests to the fungal data, only DESeq2 identified differentially abundant fungal ASVs in each of the three fields (fig. 4 & S7). This discrepancy is likely related to the sparsity of the fungal data, as ANCOM only detected structural zeroes - ASVs that were present in very few samples in at least one sample group. In Field 3, the only field where a community-level effect of disease incidence was observed, we identified eighteen fungal ASVs with differential abundance between healthy and diseased plants (fig. 4). Most of these ASVs were classified within the phylum Ascomycota and were sparsely distributed, occurring in a limited number of samples from both healthy and diseased plants. Although Fisher’s exact test suggested no significant difference in the occurrence of these ASVs between healthy and diseased plants, DESeq2 revealed significant differences in their average abundance. This highlights the utility of abundance-based methods like DESeq2 in detecting subtle but potentially biologically meaningful shifts in microbial communities.

**Figure 4.**
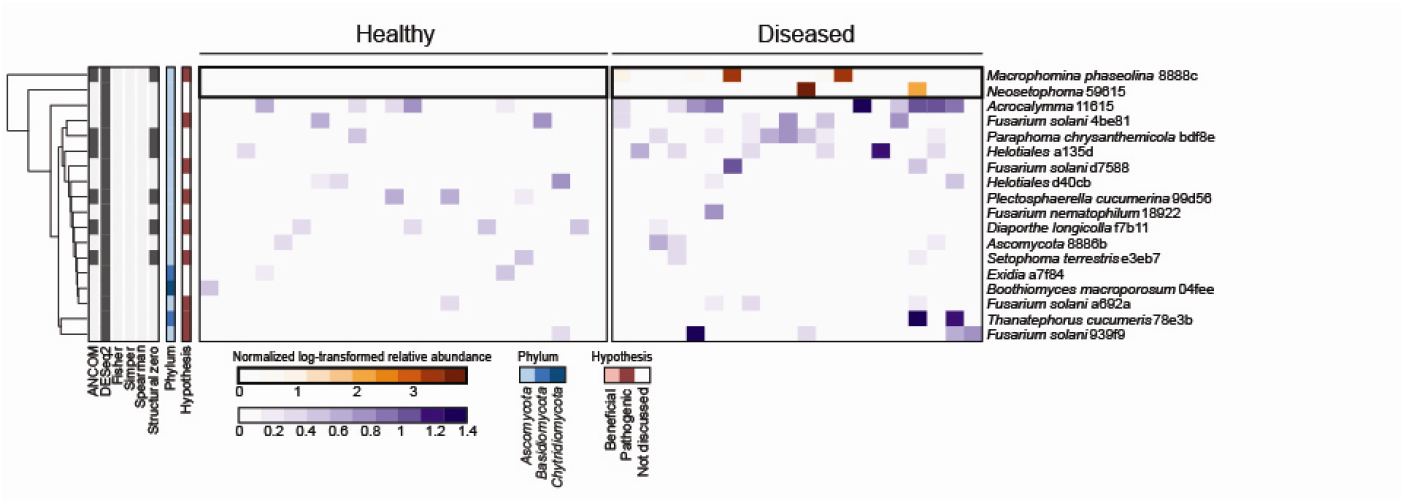
Eighteen differentially abundant fungal ASVs in the rhizosphere of healthy and naturally-infected soybean plants in Field 3. The heatmap presents log-transformed relative abundances normalized to the average relative abundance of each ASV across all healthy plant samples. Two color scales are used to enhance visualization of highly and lowly abundant ASVs. Colors to the left of the heatmap indicate whether an ASVs or its associated taxonomy is discussed in the text (red), the phylum of each ASV (blue), and the statistical method(s) by which the ASV found differentially abundant (grey). Clustering of ASVs is based on Euclidean distances.

Although we did not detect significant differences in fungal community composition between healthy and diseased plants in Fields 1 and 2, 27 fungal ASVs were differentially abundant in Field 1 and 32 ASVs in Field 2 (fig. S8). Each field included potential fungal pathogen species among the differentially abundant ASVs, such as *Fusarium solani* (Nelson, 2015), *Macrophomina phaseolina* (Mengistu *et al*., 2015; Marquez *et al*., 2021), *Diaporthe longicolla* (Li *et al*., 2015; Petrović *et al*., 2021), *Plectosphaerella cucumerina* (Hartman, 2015), *Setophoma terrestris* (Rivedal *et al*., 2022), *Thanatephorus cucumeris* (also known as *Rhizoctonia solani*; Ajayi-Oyetunde & Bradley, 2018; Yang & Hartman, 2015b), *Atractiella rhizophila* (Allen *et al*., 2020*), Cercospora* (Roy, 1982; Ward-Gauthier *et al*., 2015), *Colletotrichum* sp. (Yang & Hartman, 2015a) and *Cylindrocarpon* sp. (Mao *et al*., 2013; Tewoldemedhin *et al*., 2011). Also, ASVs representing fungal taxa with reported plant-beneficial properties were differentially abundant between healthy and diseased plants in some fields. This included *Epicoccum nigrum* (Lahlali & Hijri, 2010), *Penicillium* (Miao *et al*., 2016; Radhakrishnan *et al*., 2014), *Trichoderma strigosellum* (Pimentel *et al*., 2020) and *Clonostachys rosea* (Sun *et al*., 2020) in Field 1 and *Septoglomus viscosum* (Villani *et al*., 2021) and *C. rosea* in Field 2. However, only two fungal ASVs, *F. solani* ASV 4be81 and *M. phaseolina* ASV 8888c, were significantly enriched in the rhizosphere of diseased plants in all three fields. Remarkably, SDS-associated *Fusarium* species were not identified as differentially abundant in any field. These results show that diverse bacteria and fungi were affected in rhizosphere abundance in association with plant disease in a field-specific manner, including taxa with potential plant-beneficial or -pathogenic properties.

### *Fusarium solani* is the most likely infectious agent in the sampled soybean plants

The SDS-like disease symptoms observed in the field could be caused by several *Fusarium* spp., and 54 of the 404 unique fungal ASVs detected across the three fields were assigned to the genus *Fusarium*. These *Fusarium* ASVs were annotated as either *F. solani, Fusarium chlamydosporum* or *Fusarium nematophilum*. Thus we did not detect any of the four *Fusarium* spp. that are thought responsible for SDS in soybean. Furthermore, *F. chlamydosporum* and *F. nematophilum* have no previous connection to disease in soybean that we know of, were represented by three ASVs in total, and were present at much lower relative rhizosphere abundance than *F. solani* ASVs (fig. S9). *F. solani* was represented by 51 ASVs and the total relative abundance of these ASVs was higher in the soybean rhizosphere than in bulk soil, and higher in the rhizosphere of diseased than healthy plants (fig. 5 & S8; Wilcoxon rank sum test, adjusted *p* < 0.05). *F. solani* rhizosphere abundance did not differ between diseased plants that varied in disease severity at the plant or leaf level (ANOVA, *p* > 0.05).

**Figure 5.**
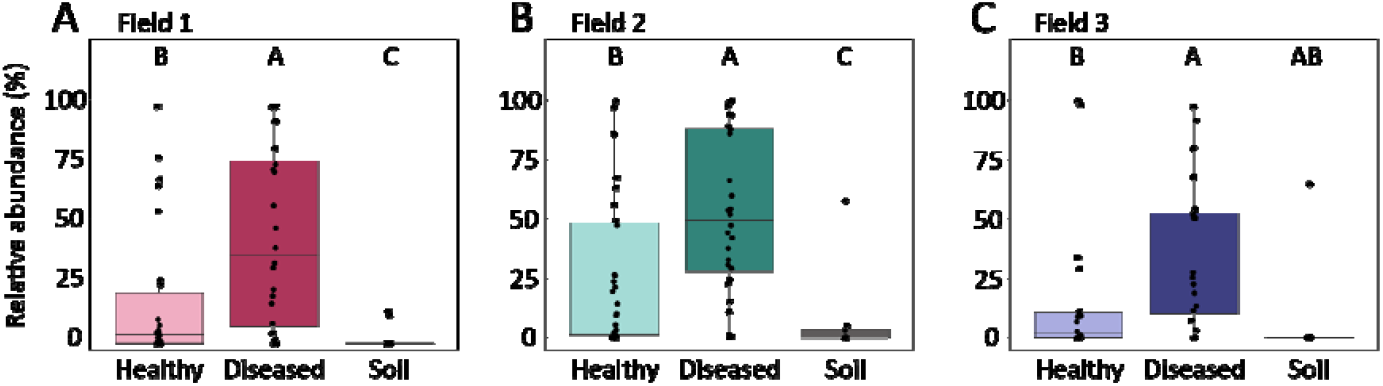
Relative abundance of *F. solani* ASVs in bulk soil and soybean rhizosphere of healthy and diseased plants. The data represents the total relative abundance of 51 unique ASVs, all annotated as *F. solani*. **A)** Field 1. Data from 26 healthy plants, 24 diseased plants and 10 soil samples. **B)** Field 2. Data from 27 healthy plants, 28 diseased plants and 8 soil samples. **C)** Field 3. Data from 22 healthy plants, 20 diseased plants and 6 soil samples. Black dots represent individual samples. Letters indicate statistically different groups based on Wilcoxon rank sum test (adjusted *p* < 0.05).

Of the 51 unique ASVs annotated as *F. solani*, 42 were detected in more than one field, with 14 found in all three fields (fig. 6A). To determine whether the three fields harboured the same *F. solani* strains, we investigated the phylogeny of the *F. solani* ASVs across the three fields by aligning their ITS2 amplicons to publicly available ITS2 sequences of *F. solani* isolates that were associated with disease in several plant species, including soybean (fig. 6B; table S2). We also included publicly available ITS2 sequences of the SDS-associated species *F. tucumaniae* and *F. virguliforme* and the ITS2 amplicons of the non-pathogenic *F. chlamydosporum* and *F. nematophilum* in our field data.

**Figure 6.**
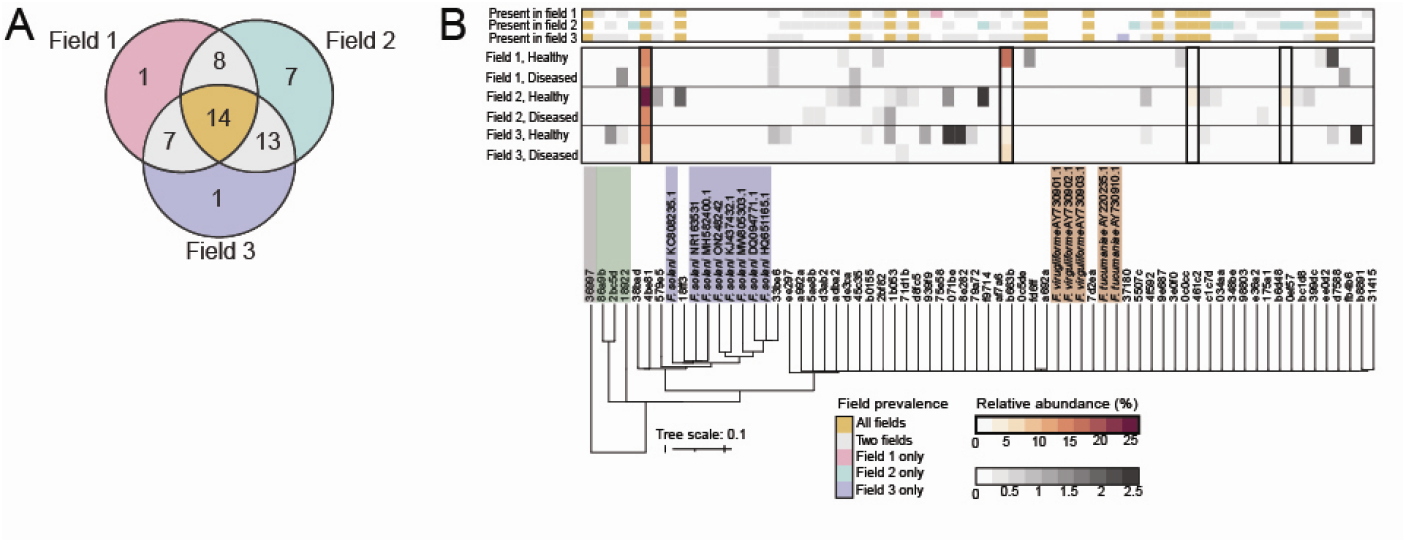
Occurrence, phylogeny and relative abundance of *Fusarium* ASVs across healthy and diseased soybean plants in three commercial fields. **A)** Venn diagram showing the number of shared and field-specific *F. solani* ASVs. **B)** Phylogenetic tree and heatmap showing the ITS2 sequence similarity and relative abundance of *F. solani* ASVs detected in rhizosphere samples from the three fields. Unique *F. solani* ASVs from the field data are identified by their five-digit hash identifiers, which distinguish them independently of taxonomic assignment. The grey label in the phylogenetic tree highlights a *Penicillium* ASV (36997), a lowly abundant ASV detected in all three fields and used as an outgroup for tree construction. A green label highlights *F. chlamydosporum* (86a9b, 2bc5d) and *F. nematophilum* (18922) ASVs. A blue label highlights publicly available ITS2 sequences of *F. solani* that have previously been isolated from diseased plants. An orange label highlights publicly available ITS2 sequences of the SDS-associated *F. tucumaniae* and *F. virguliforme*. The heatmap employs two different color scales to distinguish between highly and lowly abundant ASVs. Color annotation on the right shows the presence/absence of each *F. solani* ASV across the three fields and corresponds to the colors in A.

Three ASVs representing *F. chlamydosporum* and *F. nematophilum* cluster separately from all *F. solani* ASVs and the reference sequences of *F. solani* and the SDS-associated species *F. tucumaniae* and *F. virguliforme* (fig. 6B). Forty-six of the 51 *F. solani* ASVs were highly similar to one another and more similar to publicly available ITS2 sequences of the SDS species *F. tucumaniae* and *F. virguliforme* (99.42% average nucleotide identity; table S7) than to references sequences of *F. solani*. The UNITE database annotates these ITS2 amplicons as “species hypotheses”, where a similarity of 97-100% across the full ITS2 rRNA cistron sequence indicates likely identical species (Nilsson *et al*., 2019). Although we partially sequenced the ITS2 region, these findings indicate that the 46 ASVs may be misannotated as *F. solani* and could instead represent SDS-associated species. The relative abundance of these 46 *F. solani* ASVs is highly variable between the three soybean fields.

The remaining five ASVs annotated as *F. solani* clustered with the reference sequences of *F. solani* and thus likely represented this root rot pathogen (fig. 6B). The average percentage of shared nucleotide identity of this cluster is 96.83%, showing a slightly higher divergence among fewer ITS2 sequences than the larger cluster comprising 46 ASVs and reference sequences of SDS pathogens (fig. 6B, table S7). *F. solani* ASV 4be81 is among these 5 *F. solani* ASVs, which was consistently more abundant in the rhizosphere of diseased plants and represented the most abundant *F. solani* ASV in all fields (fig. 6B). Therefore, we hypothesize that *F. solani* ASV 4be81 is the likely causal agent of the foliar symptoms observed in the three fields.

## Discussion

Disease-induced changes in rhizosphere microbiomes have been documented in crops such as wheat, potato, and sugar beet, with several studies reporting the enrichment of disease-suppressive microbes in response to disease (Berendsen *et al*., 2018; Carrión *et al*., 2019; Liu *et al*., 2021; Yin *et al*., 2021). Field studies of soybean affected by foliar or soilborne pathogens have described differences in the rhizosphere microbiome of healthy and diseased plants though with variability in the specific taxa affected across different diseases and plant growth stages (Díaz-Cruz & Cassone, 2022; Srour *et al*., 2017). This study investigated rhizosphere microbiome changes associated with naturally-infected soybean plants across three fields in Kentucky, aiming to identify consistent disease-associated trends and potential recruitment of plant-beneficial microbes.

### Field-specific effects of disease on the rhizosphere microbiome

Plants can recruit protective microbes in response to attack (Berendsen *et al*., 2018; Goossens *et al*., 2023; Liu *et al*., 2021; Yin *et al*., 2021; Yuan *et al*., 2018). Despite similar disease severity across the three sampled fields, community-level differences between diseased and symptomless plants were only observed for bacterial communities in Field 1 and fungal communities in Field 3. Differentially abundant ASVs were detected in all fields and only two of the 62 differentially abundant fungal ASVs and one of the 50 differentially abundant bacterial ASV detected across the three fields were differentially abundant in more than a single field. The field-specific nature of these microbiome shifts reinforces the idea that soil microbial composition determines the pool of microbes available for plant-mediated recruitment in response to disease.

### High diversity of *Fusarium* species in the rhizosphere

The observed disease symptoms in the fields were initially assessed to be caused by *Fusarium* pathogens.

Remarkably, more than 10% of the detected fungal ASVs represent *Fusarium* spp., among which many ASVs were annotated as the root rot pathogen *F. solani*. Further analysis revealed that 46 of the 51 *F. solani* ASVs were misannotated, as they were highly similar to publicly available ITS2 sequences of *F. tucumaniae* and *F. virguliforme*. These *Fusarium* spp. are all considered part of the *F. solani* species complex, a group comprising more than sixty *Fusarium* spp. representing plant pathogens with a broad host range (Coleman, 2016; Geiser *et al*., 2021). However, most of the 46 ASVs belonging to the *F. solani* species complex were sparse, lowly abundant, and not consistently enriched in the rhizosphere of diseased plants, again underlining the field-specific nature of the microbiomes investigated here.

### *Fusarium solani*: the primary infectious agent

Only one Fusarium ASV, *F. solani* ASV 4be81, was relatively abundant and consistently enriched in the rhizosphere of diseased plants in all three fields. Its partial ITS2 sequence showed high similarity to sequences from *F. solani* obtained from various sources, including diseased soybean plants. This *F. solani* ASV 4be81 stood out as the most likely causal agent of the foliar chlorosis, necrosis, and premature defoliation observed in soybean plants. Only isolation of this *F. solani* strain and recapitulation of disease upon application of the isolate to healthy plants would confirm a causal relationship with the symptoms observed in the field. It should be noted that this ASV was also frequently found in plants without symptoms of disease. Perhaps symptomless plants were in premature stages of infection before symptoms became visible. Alternatively, the onset of symptoms might also rely on the presence of other rhizosphere microbes that contribute to or suppress disease.

### Potential role of *Macrophomina phaseolina* in disease development

In addition to *Fusarium* spp., other potential fungal pathogen species were detected in the sequencing data, including *M. phaseolina*, represented by a single ASV. Although this *M. phaseolina* ASV was sparsely present across the three fields, it was enriched in the rhizosphere of diseased plants. *M. phaseolina* is a generalist pathogen that can cause charcoal rot and has contributed to significant soybean losses in the United States (Allen *et al*., 2017). Although charcoal rot was not observed in the field, *F. solani* and *M. phaseolina* are known to co-infect soybean plants (Nelson, 2015), and the higher abundance *M. phaseolina* in the rhizosphere of several diseased plants across the three soybean fields might be facilitated by *F. solani* infections. Vice versa, it would be interesting to investigate whether *M. phaseolina* contributes to the development of disease by *F. solani*.

### Pathogen-driven microbiome modulation in the rhizosphere

*F. solani*, the most likely causal agent of disease in the three fields, could have reduced the impact of disease-induced, plant-driven recruitment of plant-protective microbes. Soilborne plant pathogens like *F. solani* can directly affect the abundance of soil microbes, as shown for *Verticillium dahliae* that employs an effector to inhibit the proliferation of specific soil bacteria, including pathogen-antagonistic Sphingomonads (Snelders *et al*., 2020). Certain soil microbes can also facilitate pathogen infection (Dewey *et al*., 1999; Li *et al*., 2019) and thus may be supported by plant pathogens. Such pathogen-driven microbiome modulation could obscure plant-induced processes, such as the recruitment of plant-beneficial microbes, and thereby reduce the detectable changes in the abundance of specific microbial features across multiple fields.

### Taxa enriched on healthy plants may have prevented disease

We identified differentially abundant ASVs in each field that belonged to microbial taxa with reported plant-beneficial properties. This included ASVs representing the fungal species *C. rosea, Penicillium* and *Trichoderma*, that have previously been shown to inhibit *Fusarium* growth *in vitro* or enhance plant resistance against *Fusarium* infection (Miao *et al*., 2016; Pimentel *et al*., 2020; Radhakrishnan *et al*., 2014; Sun *et al*., 2020). Several ASVs representing these three fungal taxa were more abundant in the rhizosphere of healthy plants compared to diseased plants. These fungi were thus not recruited by the plant upon pathogen infection; however, they could have protected healthy soybean plants against infection by *F. solani*. Initial microbiome communities can be predictive of disease onset, as has been shown by Gu *et al*. (2022) and Wei *et al*. (2019).

### Recruitment of *Sphingomonas* in diseased plants

Only a single bacterial ASV was enriched in the rhizosphere of diseased soybean plants in two fields; a *Sphingomonas* ASV that might represent a microbe recruited by soybean upon pathogen infection. Members of the *Sphingomonas* genus have been connected to disease suppression of several bacterial and fungal pathogens in other plant species (Innerebner *et al*., 2011; Kyselková *et al*., 2014; Wei *et al*., 2019), and the soilborne pathogen *Verticillium dahliae* has evolved an effector to specifically suppress antagonistic Sphingomonads (Snelders *et al*., 2020). These findings underline the relevance of Sphingomonads in disease ecology, and it would be interesting to investigate whether such a relationship could play a role in the suppression of *F. solani*-induced disease in soybean.

### Conclusion: Microbial interplay in soybean disease ecology

This study highlights the complexity of disease ecology, showing field-specific microbiome differences between healthy and naturally-infected plants. Several candidate plant-beneficial taxa were differentially abundant between healthy and diseased plants, warranting further study for potential application in crop protection. We identified *Fusarium solani* as the likely disease-causing agent, potentially co-infecting plants with the root pathogen *M. phaseolina*. Our findings emphasize the interplay between plant pathogens, candidate plant-beneficial microbes, and field-specific microbiomes, underscoring the importance of the entire microbial rhizosphere community in disease suppression and plant health.

## Acknowledgements

The authors thank the soybean growers who granted permission to sample their fields. Special thanks go to Andrew Gordon and Koppert Biological Systems for their extensive help in setting up and carrying out the field sampling campaign.

## Contributions

**S.v.B**.: methodology, investigation, formal analysis, data curation, writing – original draft, visualization, funding acquisition. **B.S.O**., **A.D.G**.: methodology, investigation, writing – review & editing. **H.A.v.P**.: methodology. **P.A.H.M.B**.: conceptualization, methodology, supervision, writing – review & editing. **C.M.J.P**.: conceptualization, methodology, supervision, writing – review & editing. **S.L.L**.: methodology, investigation, writing – review & editing. **R.L.B**.: conceptualization, methodology, supervision, writing – review & editing, funding acquisition.

