## Supplemental information for "Microbiome responses to natural *Fusarium* infection in field-grown soybean plants"

**Supplemental figures**

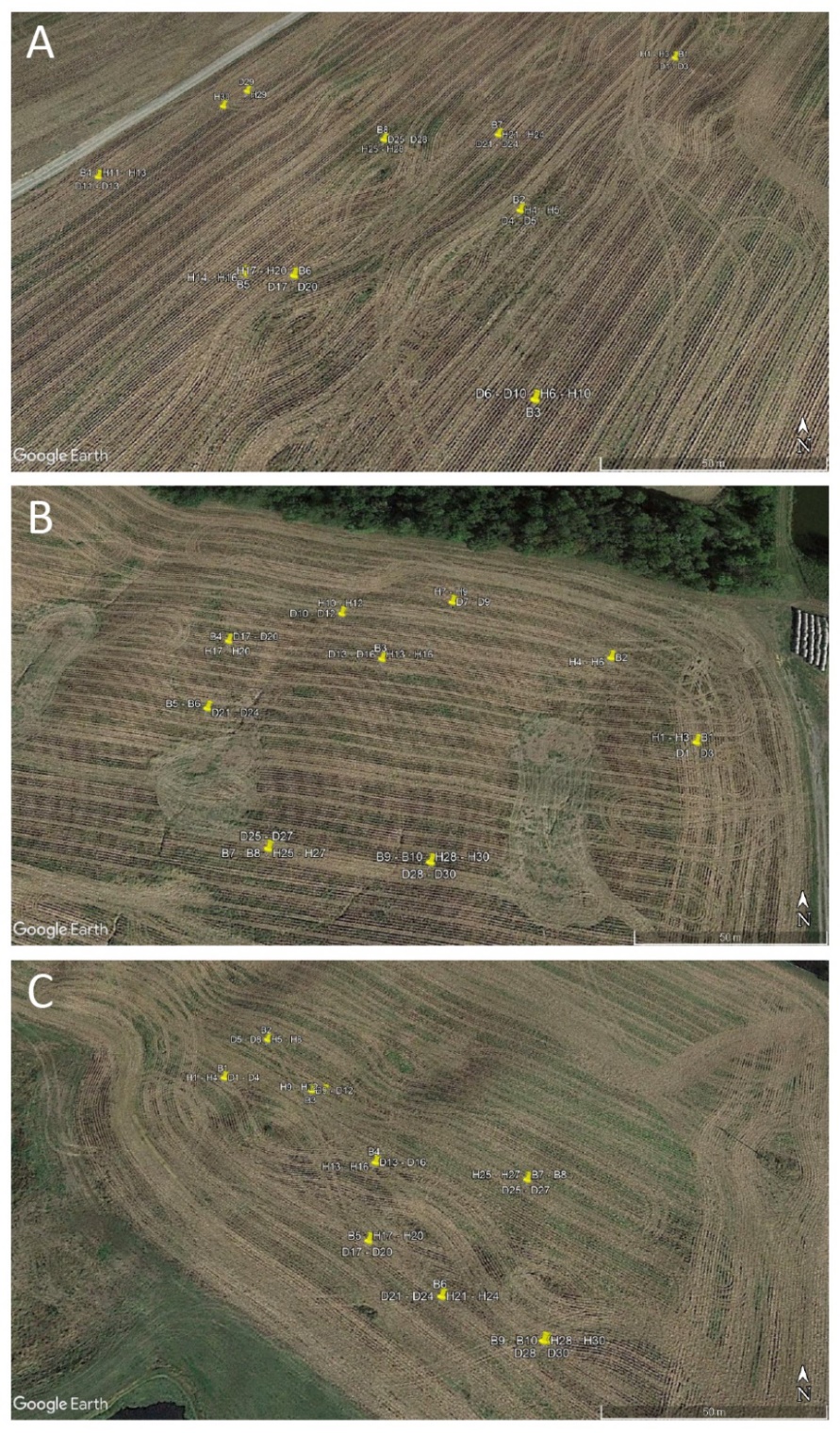

**Figure S1. Location of soybean plants and bulk soil sampled in three soybean fields.** **A)** Sample locations in Field 1. Samples D14-D16 could not be plotted but were close to B5 and H14-H16. Similarly, B9 and B10 were close to H29 and D30. **B)** Sample locations in Field 2. Samples D4-D6 could not be plotted but were close to B2 and H4-H6. H21-H24 were close to B5-B6 and D21-D24. **C)** Sample locations in Field 3. GPS coordinates were recorded for each plant and soil sample and plotted in Google Earth Pro. B = bulk soil; H = healthy plant; D = diseased plant. Numbers represent sample number up to a total of ten bulk soil and thirty healthy/diseased plant samples per field.

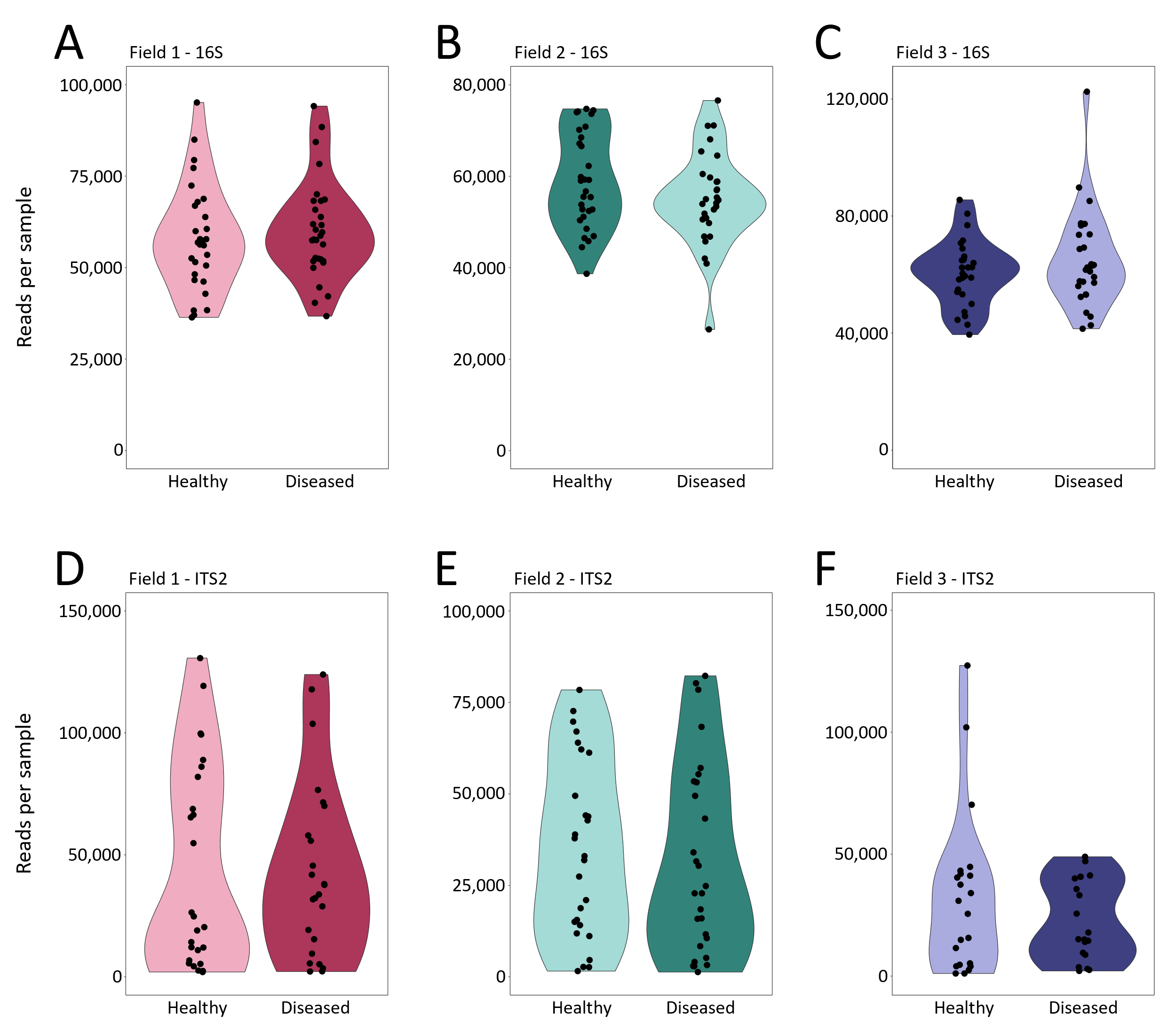

**Figure S2.** **The average read depth of 16S and ITS2 amplicons per sample of healthy and diseased plants across three fields.** **A)** Field 1, 16S. **B)** Field 2, 16S. **C)** Field 3, 16S. **D)** Field 1, ITS2. **E)** Field 2, ITS2. **F)** Field 3, ITS2.

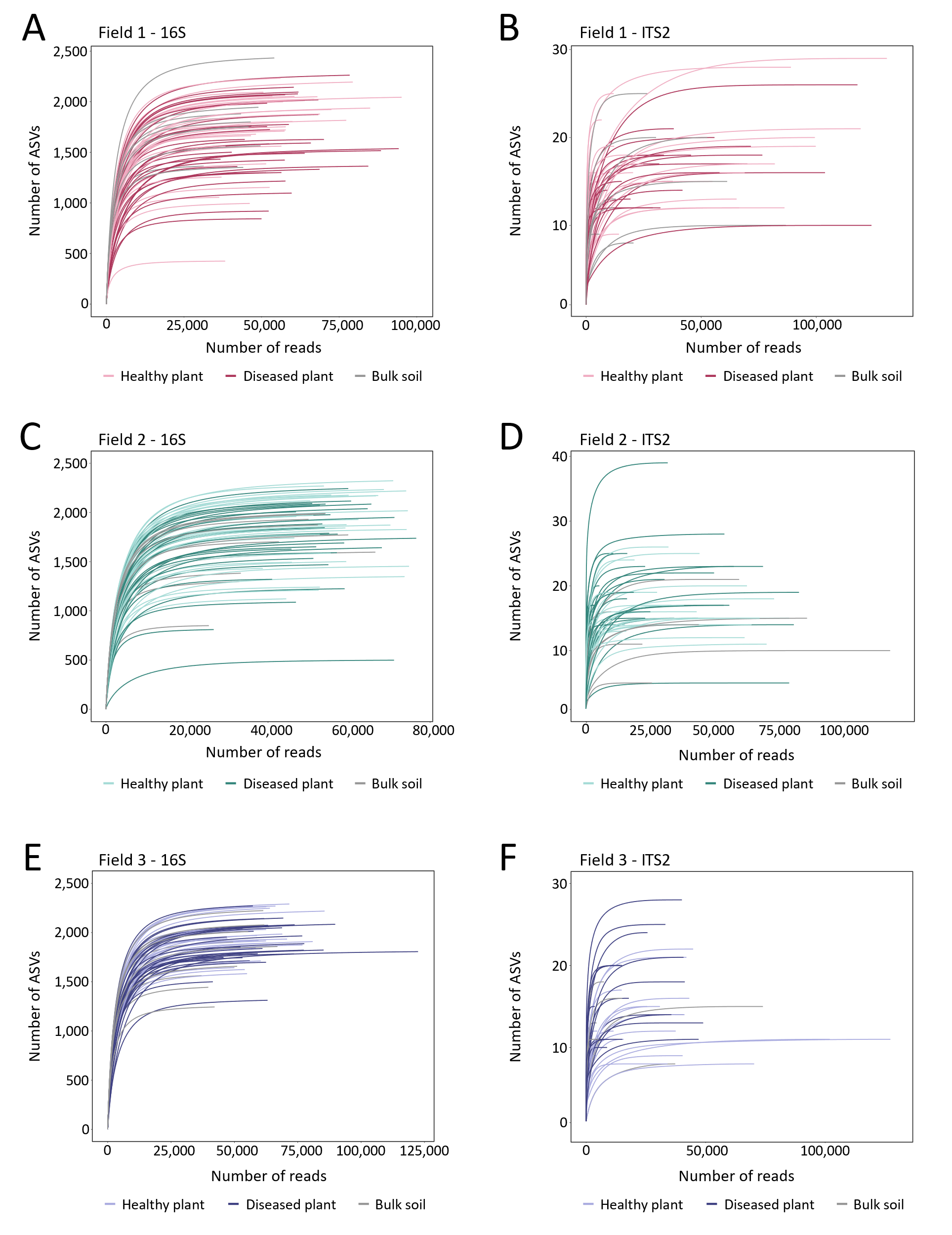

**Figure S3. The number of 16S and ITS2 ASVs detected per sample with increasing sequencing depth in healthy plant rhizosphere, diseased plant rhizosphere and bulk soil across three fields.** **A)** Field 1, 16S2. **B)** Field 1, ITS2. **C)** Field 2, 16S. **D)** Field 2, ITS2. **E)** Field 3, 16S. **F)** Field 3, ITS2.

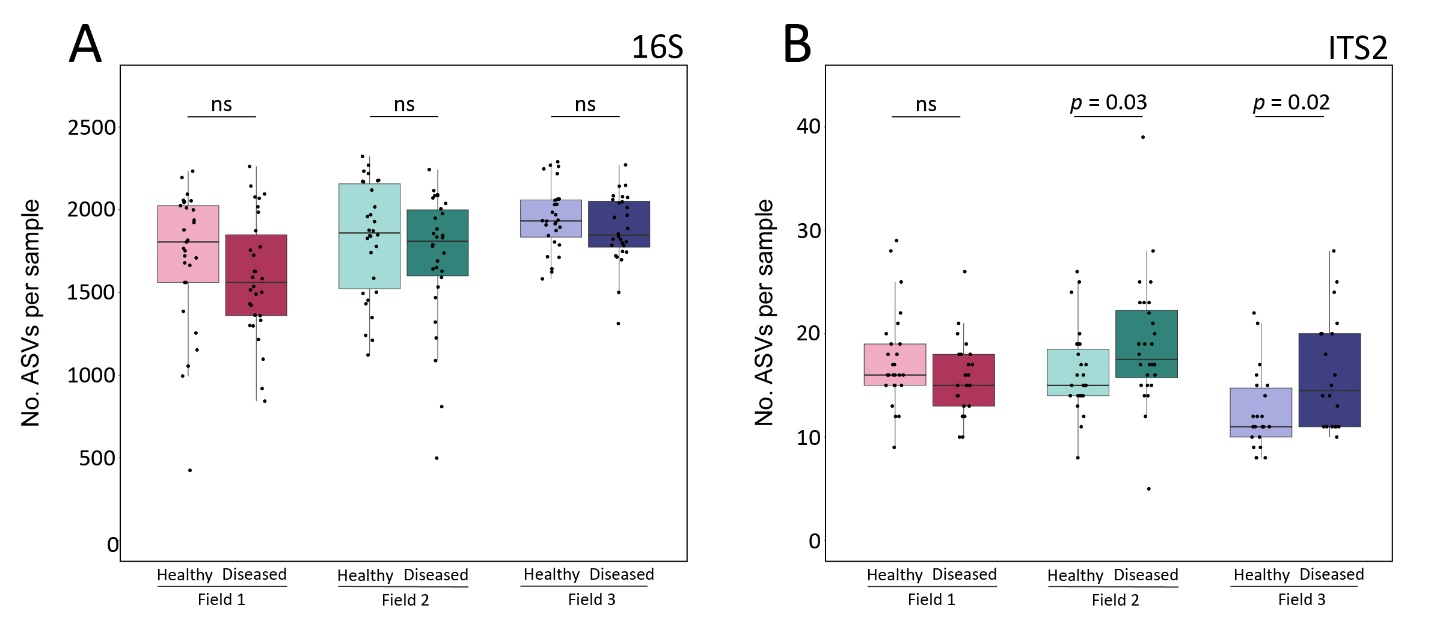

**Figure S4. The number of 16S and ITS2 ASVs detected per rhizosphere sample of healthy or diseased soybean plants. A)** 16S data. **B)** ITS2 data. Each dot represents the total number of ASVs found in a single rhizosphere sample.

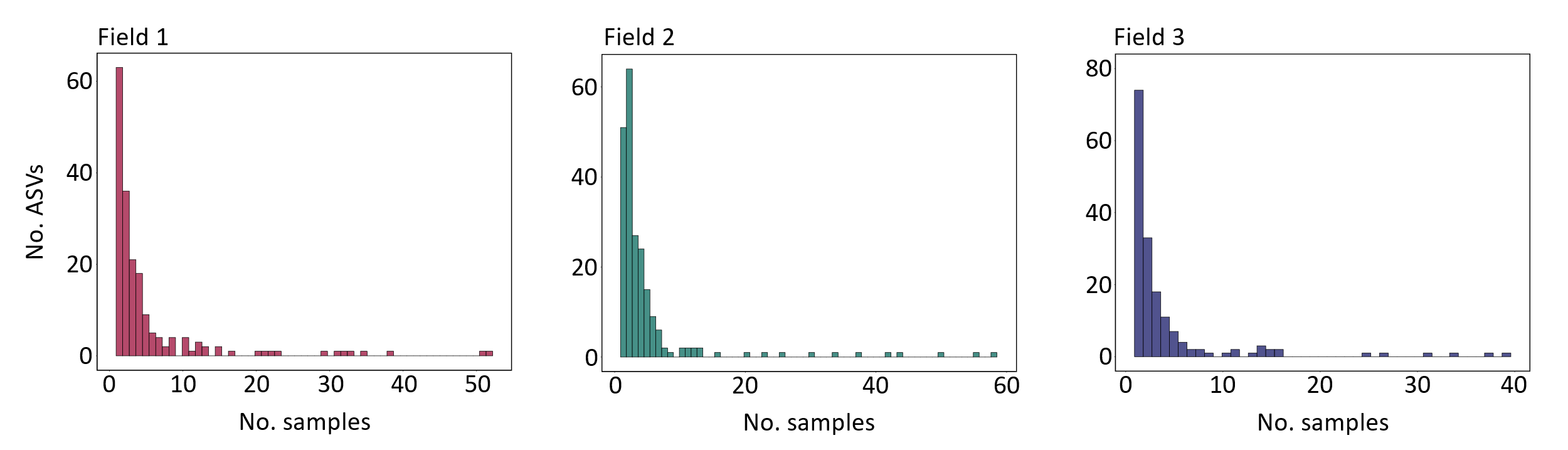

**Figure S5.** **The occupancy of ITS2 ASVs across rhizosphere and bulk soil samples from three soybean fields.**

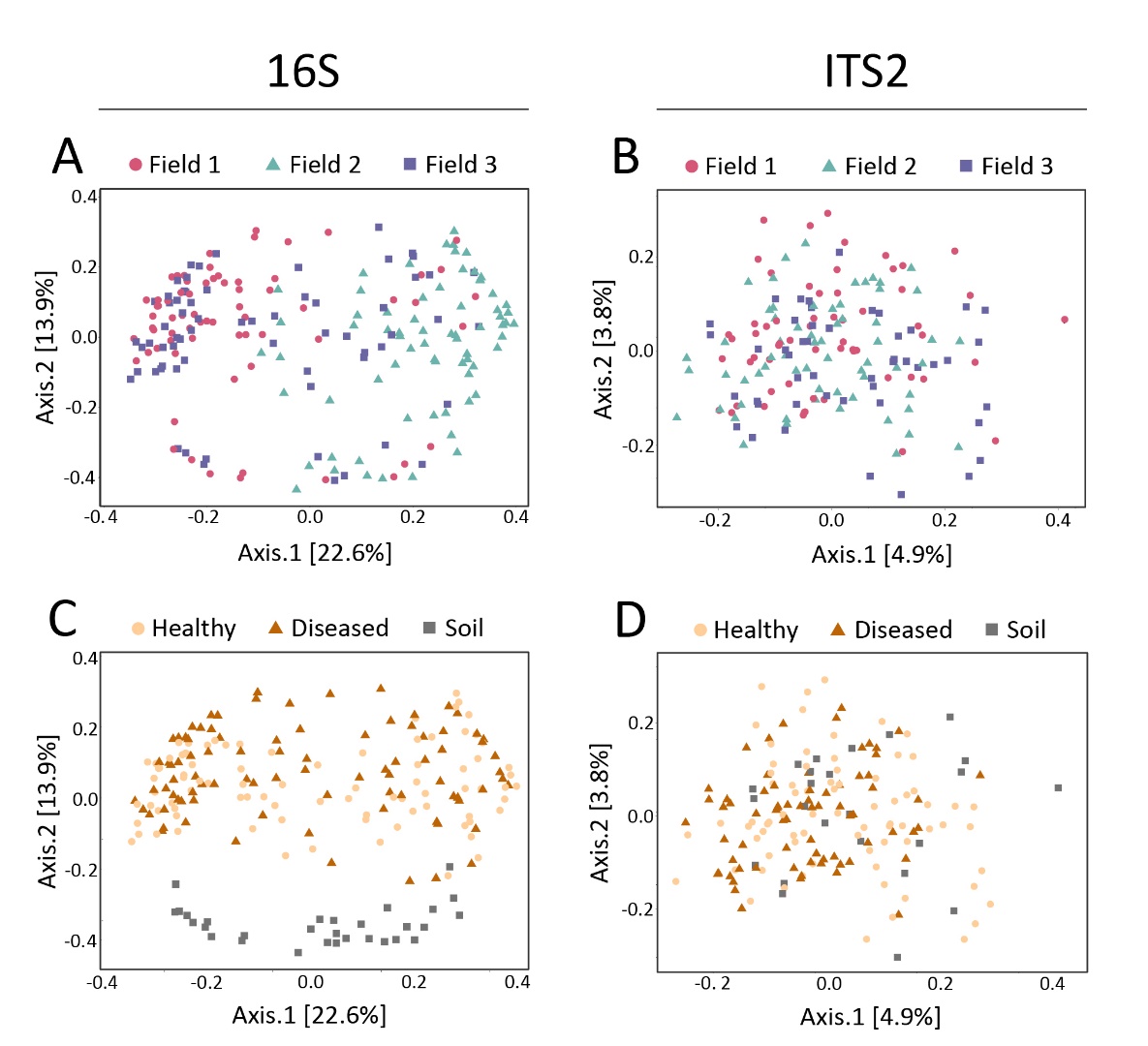

**Figure S6. Principal coordinate analysis (PCoA) plots of bacterial and fungal communities of soybean rhizosphere and bulk soil samples from three fields. A)** PCoA of bacterial communities based on Bray-Curtis distances, combining rhizosphere and bulk soil samples, colored by field of sampling. **B)** PCoA of fungal communities based on Jaccard distances, combining rhizosphere and bulk soil samples, colored by field of sampling. **C)** Same plot as A, but colored by sample type. **D)** Same plot as B, but colored by sample type. Healthy = healthy plant rhizosphere; Diseased = diseased plant rhizosphere; Soil = bulk soil. Statistics describing differences between bacterial and fungal communities of healthy plant rhizosphere, diseased plant rhizosphere and bulk soil are provided in tables S1 & S2.

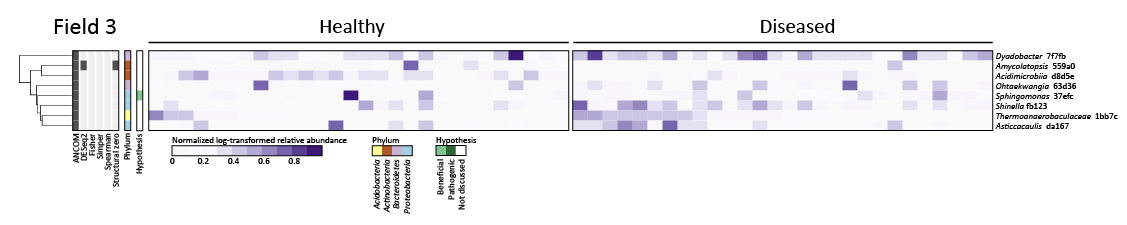

**Figure S7. Heatmap of differentially abundant bacterial ASVs in the rhizosphere of healthy and diseased soybean plants in Field 3.** The heatmap shows the log-transformed relative abundances normalized by the average relative abundance of that ASV across all healthy plant samples. Color annotation on the left of the heatmap indicates phylum annotation of each ASV; grey annotation indicates statistical method(s) by which each differentially abundant ASV was identified; tree branches indicate clustering of the normalized relative abundance of ASVs based on Euclidean distance measures.

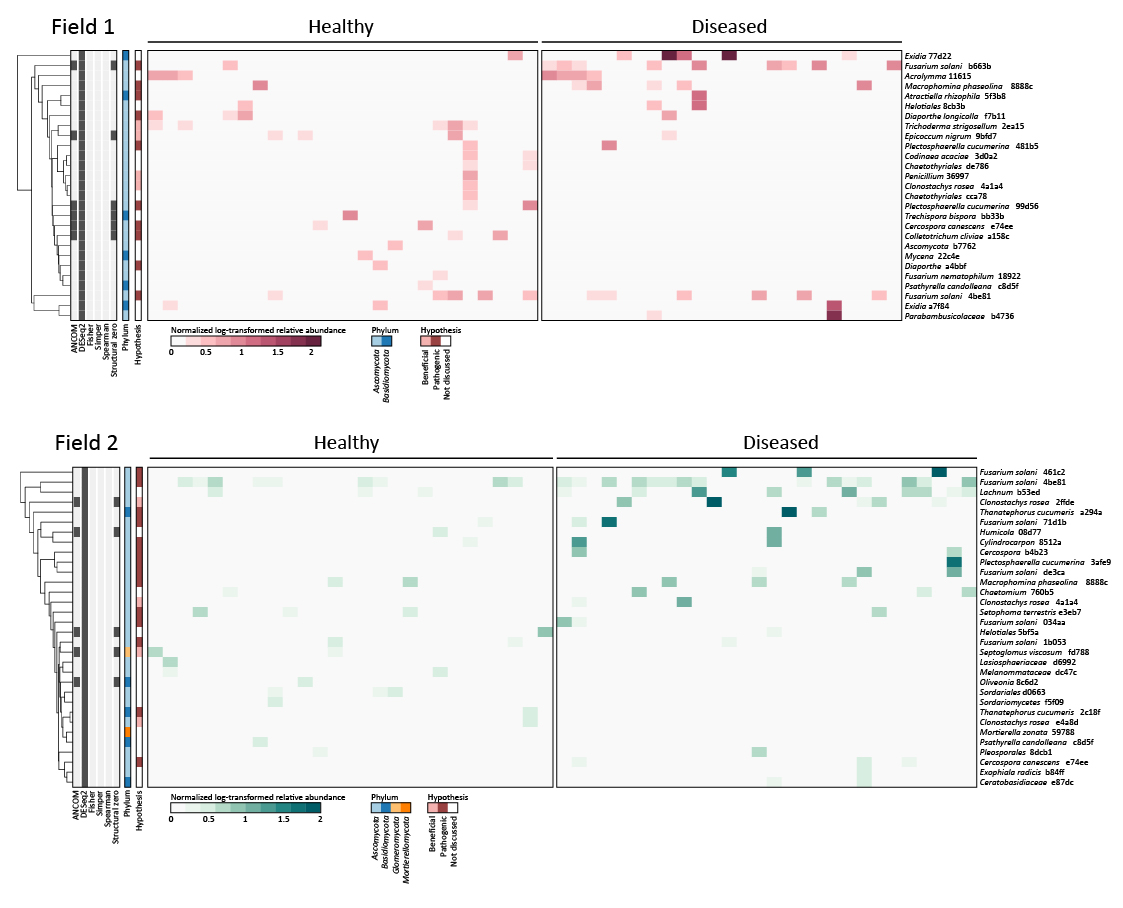
**Figure S8. Heatmaps of differentially abundant fungal ASVs in the rhizosphere of healthy and diseased soybean plants in Field 1 (top) and Field 2 (bottom).** The heatmaps show the log-transformed relative abundances normalized by the average relative abundance of that ASV across all healthy plant samples. Color annotation on the left of each heatmap indicates phylum annotation of each ASV; grey annotation indicates statistical method(s) by which each differentially abundant ASV was identified; tree branches indicate clustering of the normalized relative abundance of ASVs based on Euclidean distance measures.

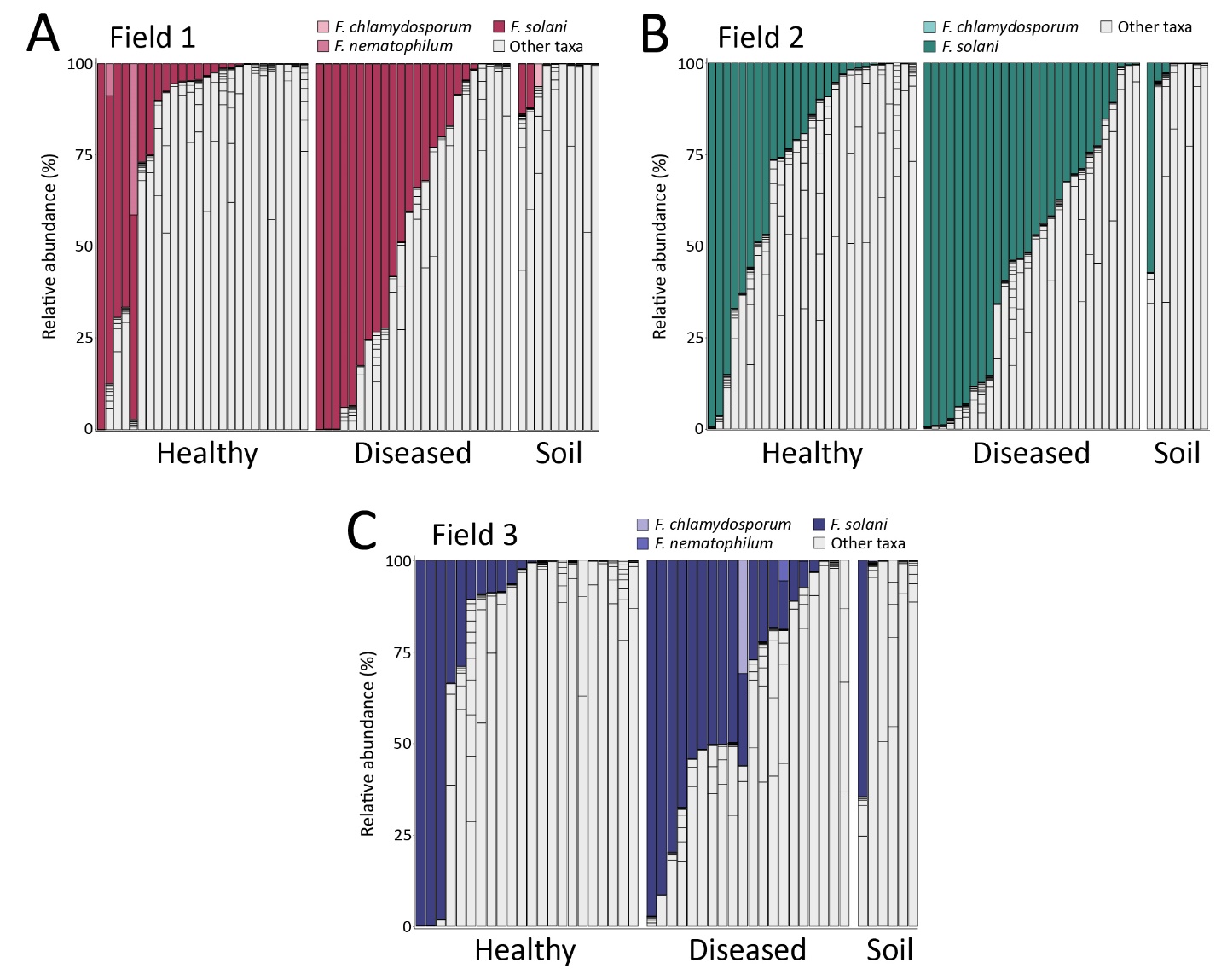
 **Figure S9.** **The relative abundance of ITS2 ASVs annotated as *F. solani* in rhizosphere and soil samples of three commercial soybean fields.** **A)** *F. solani* relative abundance in Field 1. **B)** *F. solani* relative abundance in Field 2. **C.** *F. solani* relative abundance in Field 3. Each bar represents one sample.

**Supplemental tables**

**Table S1. Primer sets used to generate 16S and ITS2 amplicons of soybean rhizosphere and bulk soil samples of three commercial fields.**

| **Name** | **Amplicon** | **Primer Type** | **Forward/**  **Reverse** | **Reference** | **Sequence** |
| --- | --- | --- | --- | --- | --- |
| ITS2-fw | ITS2 | Amplicon primer | Forward | Génome Quebec (CN) | ACACTCTTTCCCTACACGACGCTCTTCCGATCTGAACGCAGCRAAIIGYGA |
| ITS2-rv | ITS2 | Amplicon primer | Reverse | Génome Quebec (CN) | GTGACTGGAGTTCAGACGTGTGCTCTTCCGATCTTCCTCCGCTTATTGATATGC |
| 341FP1-FwR1 | 16S | Amplicon primer | Forward | Génome Quebec (CN) | ACACTCTTTCCCTACACGACGCTCTTCCGATCTCCTACGGGNGGCWGCAG |
| 341FP2-FwR1 | 16S | Amplicon primer | Forward | Génome Quebec (CN) | ACACTCTTTCCCTACACGACGCTCTTCCGATCTTCCTACGGGNGGCWGCAG |
| 341FP3-FwR1 | 16S | Amplicon primer | Forward | Génome Quebec (CN) | ACACTCTTTCCCTACACGACGCTCTTCCGATCTACCCTACGGGNGGCWGCAG |
| 341FP4-FwR1 | 16S | Amplicon primer | Forward | Génome Quebec (CN) | ACACTCTTTCCCTACACGACGCTCTTCCGATCTCAACCTACGGGNGGCWGCAG |
| 805RP1-RvR2 | 16S | Amplicon primer | Reverse | Génome Quebec (CN) | GTGACTGGAGTTCAGACGTGTGCTCTTCCGATCTGACTACHVGGGTATCTAATCC |
| 805RP2-RvR2 | 16S | Amplicon primer | Reverse | Génome Quebec (CN) | GTGACTGGAGTTCAGACGTGTGCTCTTCCGATCTTGACTACHVGGGTATCTAATCC |
| 805RP3-RvR2 | 16S | Amplicon primer | Reverse | Génome Quebec (CN) | GTGACTGGAGTTCAGACGTGTGCTCTTCCGATCTACGACTACHVGGGTATCTAATCC |
| 805RP4-RvR2 | 16S | Amplicon primer | Reverse | Génome Quebec (CN) | GTGACTGGAGTTCAGACGTGTGCTCTTCCGATCTCATGACTACHVGGGTATCTAATCC |
| ITS2-block-fw | ITS2 | Blocking primer | NA | Agler *et al.*, 2016 (BioRXiv) | CGTCTGCCTGGGTGTCACAAATCGTCGTCC |
| ITS2-block-rv | ITS2 | Blocking primer | NA | Agler *et al.*, 2016 (BioRXiv) | CCTGGTGTCGCTATATGGACTTTGGGTCAT |
| mPNA | 16S | Blocking primer | NA | Lundberg *et al.*, 2013 | GGCAAGTGTTCTTCGGA |
| pPNA | 16S | Blocking primer | NA | Lundberg *et al.*, 2013 | GGCTCAACCCTGGACAG |

**Table S2. *Fusarium* ITS2 sequences obtained from the NCBI GenBank database to align with ITS2 sequences obtained from soybean rhizosphere samples.**

| **ID** | **Source** | **NCBI identifier** | **Reference** |
| --- | --- | --- | --- |
| *Fusarium solani* DQ094771.1 | Red clover | DQ094771.1 | Zhang *et al.,*2006 |
| *Fusarium solani* HQ651165.1 | *Grevillea robusta* | HQ651165.1 | Njuguna *et al.,*2010 |
| *Fusarium solani* KC808235.1 | Unknown | KC808235.1 | O'Donnell *et al.,*2013 |
| *Fusarium solani* KJ437432.1 | Soybean | KJ437432.1 | Chitrampalam & Nelson Jr, 2015 |
| *Fusarium solani* MH582400.1 | River sediment | MH582400.1 | O'Donnell *et al.,*2018 |
| *Fusarium solani* MW805303.1 | Unknown | MW805303.1 | Zuo *et al.,*2021 |
| *Fusarium solani* NR_163531 | Potato tuber | NR_163531 | Schroers *et al.,*2016 |
| *Fusarium solani* ON248242 | Unknown | ON248242 | Wang & Wang, 2022 |
| *Fusarium tucumaniae* AY220235.1 | Soybean | AY220235.1 | Aoki *et al.,*2003 |
| *Fusarium tucumaniae* AY730910.1 | Soybean | AY730910.1 | Aoki *et al.,*2005 |
| *Fusarium virguliforme* AY730901.1 | Soybean | AY730901.1 | Aoki *et al.,*2005 |
| *Fusarium virguliforme* AY730902.1 | Soybean | AY730902.1 | Aoki *et al.,*2005 |
| *Fusarium virguliforme* AY730903.1 | Soybean | AY730903.1 | Aoki *et al.,*2005 |

**Table S3. Pairwise statistical comparisons by PERMANOVA of bacterial communities between fields (1-3) and sample type (healthy plant rhizosphere, diseased plant rhizosphere, bulk soil).** Comparisons were performed for the Bray-Curtis distance metric. The R^2^ and FDR-corrected *p*-values for each comparison are reported.

| **Field** | Field 1 | Field 2 | Field 3 |
| --- | --- | --- | --- |
| Field 1 |  | R^2^ = 0.15 | R^2^ = 0.03 |
| Field 2 | Adjusted *p* < 0.01 |  | R^2^ = 0.14 |
| Field 3 | Adjusted *p* < 0.01 | Adjusted *p* < 0.01 |  |
| **Sample Type** | Healthy | Diseased | Bulk soil |
| Healthy |  | R^2^ = 0.01 | R^2^ = 0.15 |
| Diseased | Adjusted *p* = 0.06 |  | R^2^ = 0.16 |
| Bulk soil | Adjusted *p* < 0.01 | Adjusted *p* < 0.01 |  |

**Table S4. Pairwise statistical comparisons by PERMANOVA of fungal communities between fields (1-3) and sample type (healthy plant rhizosphere, diseased plant rhizosphere, bulk soil).** Comparisons were performed for the Jaccard distance metric. The R^2^ and FDR-corrected *p*-values for each comparison are reported.

| **Field** | Field 1 | Field 2 | Field 3 |
| --- | --- | --- | --- |
| Field 1 |  | R^2^ = 0.02 | R^2^ = 0.02 |
| Field 2 | Adjusted *p* < 0.01 |  | R^2^ = 0.02 |
| Field 3 | Adjusted *p* < 0.01 | Adjusted *p* < 0.01 |  |
| **Sample Type** | Healthy | Diseased | Bulk soil |
| Healthy |  | R^2^ = 0.01 | R^2^ = 0.01 |
| Diseased | Adjusted *p* = 0.03 |  | R^2^ = 0.02 |
| Bulk soil | Adjusted *p* = 0.58 | Adjusted *p* < 0.01 |  |

**Table S5. Pairwise statistical comparisons by PERMANOVA of bacterial communities in the rhizosphere of healthy plants, the rhizosphere of diseased plants and bulk soil in each of three commercial soybean fields.** Comparisons were performed for the Bray-Curtis distance metric. The R^2^ and FDR-corrected *p*-values for each comparison are reported.

| **Field 1** | Healthy | Diseased | Bulk soil |
| --- | --- | --- | --- |
| Healthy |  | R^2^ = 0.04 | R^2^ = 0.22 |
| Diseased | Adjusted *p* = 0.01 |  | R^2^ = 0.22 |
| Bulk soil | Adjusted *p* < 0.01 | Adjusted *p* < 0.01 |  |
| **Field 2** | Healthy | Diseased | Bulk soil |
| Healthy |  | R^2^ = 0.01 | R^2^ = 0.23 |
| Diseased | Adjusted *p* = 0.68 |  | R^2^ = 0.21 |
| Bulk soil | Adjusted *p* < 0.01 | Adjusted *p* < 0.01 |  |
| **Field 3** | Healthy | Diseased | Bulk soil |
| Healthy |  | R^2^ = 0.02 | R^2^ = 0.17 |
| Diseased | Adjusted *p* = 0.30 |  | R^2^ = 0.20 |
| Bulk soil | Adjusted *p* < 0.01 | Adjusted *p* < 0.01 |  |

**Table S6. Pairwise statistical comparisons by PERMANOVA of fungal communities in the rhizosphere of healthy plants, the rhizosphere of diseased plants and bulk soil in each of three commercial soybean fields.** Comparisons were performed for the Jaccard distance metric. The R^2^ and FDR-corrected *p*-values for each comparison are reported.

| **Field 1** | Healthy | Diseased | Bulk soil |
| --- | --- | --- | --- |
| Healthy |  | R^2^ = 0.02 | R^2^ = 0.03 |
| Diseased | Adjusted *p* = 1.00 |  | R^2^ = 0.04 |
| Bulk soil | Adjusted *p* = 0.38 | Adjusted *p* = 0.04 |  |
| **Field 2** | Healthy | Diseased | Bulk soil |
| Healthy |  | R^2^ = 0.02 | R^2^ = 0.03 |
| Diseased | Adjusted *p* = 0.54 |  | R^2^ = 0.05 |
| Bulk soil | Adjusted *p* = 0.53 | Adjusted *p* = 0.02 |  |
| **Field 3** | Healthy | Diseased | Bulk soil |
| Healthy |  | R^2^ = 0.04 | R^2^ = 0.04 |
| Diseased | Adjusted *p* < 0.01 |  | R^2^ = 0.04 |
| Bulk soil | Adjusted *p* = 1.00 | Adjusted *p* = 1.00 |  |

**Table S7. Percent identity values for ITS2 sequences aligned with the MAFFT tool of EMBL-EBI.** A color range from red to green indicates relatively low and high percent identity values, respectively. Three groups of sequences are indicated with thick cell borders and colors in rows and columns. Grey column/row annotation indicates the smaller cluster of *F. solani* ASVs clustering with *F. solani* reference sequences in the heatmap of fig. 3. Blue column/row annotation indicates the large cluster of 51 *F. solani* ASVs and reference sequences for the SDS pathogens *F. tucumaniae* and *F. virguliforme* in fig. 3. Due to the large size of this table, a QR-code to the online open repository Zenodo is presented here.

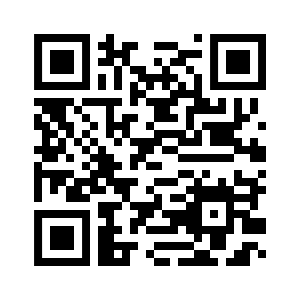

**Supplemental Methods**

**DNA extractions from soybean rhizosphere and bulk soil samples**

This protocol was used to extract DNA from soybean rhizosphere (soil attached to roots) and bulk soil samples. The procedures are based on the DNeasy PowerSoil kit (Qiagen, Hilden, Germany) and the patent of its predecessor, the MoBio PowerSoil DNA isolation kit. These kits were designed for small soil samples (up to 0.25g) and their protocols were adjusted to enable processing of larger soybean root samples.

This protocol makes use of the KingFisher Flex Purification System (ThermoFisher), designed for high-throughput reactions in 96-well plates.

Recipes for chemical solutions can be found at the end of this protocol.

Note: all samples (soil, roots) were frozen or kept on ice at all times prior to cell lysis (step 2). For steps with smaller volumes, use 2-ml Eppendorf tubes (not 1.5-ml tubes).

Step-by-step procedures

1) Determine the weight of the (root) samples. Make sure that samples do not thaw during this process.

2) In preparation of cell lysis, calculate and prepare the volume of bead solution and C1 solution to be added to each sample. For 1 g of sample, prepare 3 ml bead solution and 0.24 ml C1 solution. Make sure that the pH of the bead solution is adjusted to 8.0-8.5. Be aware that the total volume of lysis solution should be sufficient to transfer supernatant to a clean tube (e.g. in 50-ml Falcon tubes, we required a total volume of ≥ 11 ml for each soybean root sample).

3) Perform cell lysis with a mixture of bead solution and C1 solution. Add 3 ml bead solution and 0.24 ml C1 solution per gram of sample. Invert samples several times by hand and place samples in a 70°C water bath for 10 min.

4) Shake the samples with sterile 3-mm glass beads (10 beads for soybean roots, 2 beads for soil samples) in a Tissuelyser II (2-ml tubes; Qiagen, Hilden, Germany) or paint shaker (50-ml tubes; SK550 1.1 heavy-duty paint shaker, Fast & Fluid, Sassenheim, the Netherlands). Set Tissuelyser twice for 10 min at 30 Hz, or the paint shaker at 270 sec at speed 3, 270 sec at speed 6. Note: soybean roots were not fully grinded during this process.

5) Spin down suspended particles (max 10.000g). For 50-ml tubes, centrifuging at 4,816 g for 2 min in a swing-out centrifuge should be sufficient.

6) Transfer 700 μl supernatant from each sample to clean 2.0-ml tubes.

7) Clean up the supernatant with 250 μl C2 solution (Qiagen, Hilden, Germany). Vortex for 5 seconds and put samples on ice for 5 min. Centrifuge samples for 1 min at 10.000 rcf. Transfer 600 μl supernatant to a clean tube.

8) Add 200 μl C3 solution (Qiagen, Hilden, Germany) to the new tubes. Vortex for 3 seconds and put on ice for 5 min. Centrifuge samples for 1 min at 10.000g. Transfer up to 750 μl to a clean tube.

9) For 96 samples, prepare:

• a binding bead mix of 45 ml binding solution (ThermoFisher) and 2 ml MagMax beads (ThermoFisher). Be aware that the beads are viscous and need to be pipetted with care.

• 1 Kingfisher deep-well plate with 500 μl sample with 500 μl binding bead mix per well (plate ‘Bind’). Make sure that the MagMax beads are suspended properly in the binding solution before pipetting into the plate.

• 1 Kingfisher microplate with 50 μl C6 solution per well (plate ‘Elution’).

• 2 Kingfisher deep-well plates with 1000 μl C5 solution per well (plates ‘Wash 1’ and ‘Wash 2’).

• 2 Kingfisher deep-well plates with 1000 μl 80% ethanol per well (plates ‘Wash 3’ and ‘Wash 4’).

• 1 Kingfisher deep-well plate with a 96-tip comb (plate ‘Tip’).

10) Set up computer-steered protocol for the KingFisher Flex Purification System (ThermoFisher), see detailed protocol below. Start protocol and load plates in correct order as prompted. Start protocol (takes approx. 1 hour).

11) Check DNA quantities and overall quality with Nanodrop (ThermoFisher, Waltham, United States). Note that the amounts of DNA extracted from soil samples with this protocol can seem very low but these samples can often still be amplified by PCR. If you require double-stranded DNA, (also) check DNA quantities with a fluorescent dye in a system such as Qubit (Invitrogen, Waltham, United States).

Recipes for DNA extraction solutions

Bead solution (1L):

NaH_2_PO_4_ (362 mM)	500 ml

Gu thiocyanate (CH_6_N_3_; 605 mM)	200 ml

MQ	300 ml

pH adjusted to 8.0-8.5 with NaOH (note: use high concentration NaOH, needs many drops)

C1 solution (100 ml):

NaCl (1.5M)	10 ml

SDS (8%)	50 ml

Tris (1.5 M)	33.33 ml

MQ	15.67 ml

C5 solution (20 ml):

Tris (1.5M)	0.1 ml

NaCl (1.5M)	1.3 ml

EtOH (96%)	10.4 ml

MQ	8.1 ml

C6 solution (1 ml):

Tris (100 mM, pH adjusted to 8.0-8.5)	0.1 ml

MQ	0.9 ml

**Detailed protocol for DNA extraction with Kingfisher Flex Purification System**

The screenshot below shows the overview of steps performed with the Kingfisher Flex Purification System (ThermoFisher). In each step, a plate prepared in the procedures above is selected for action:

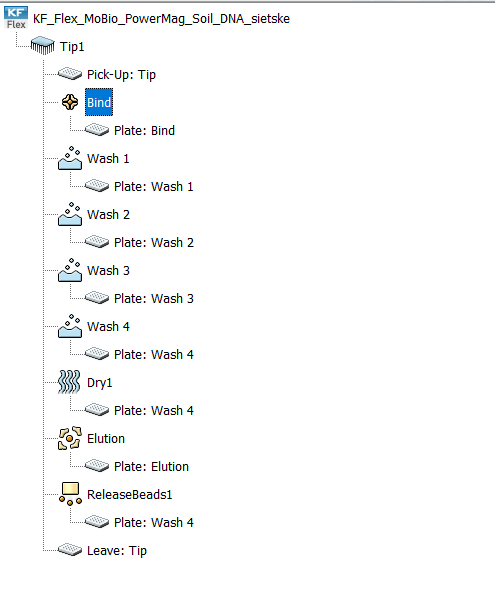

In the BindIt software for KingFisher machines, create the protocol as follows:

1) Add the ‘Pick-Up’ step (select plate ‘Tip’).

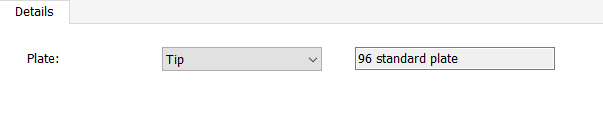

2) Add a ‘Bind’ step where samples and beads are mixed, and beads are subsequently bound to tips (select plate ‘Bind’).

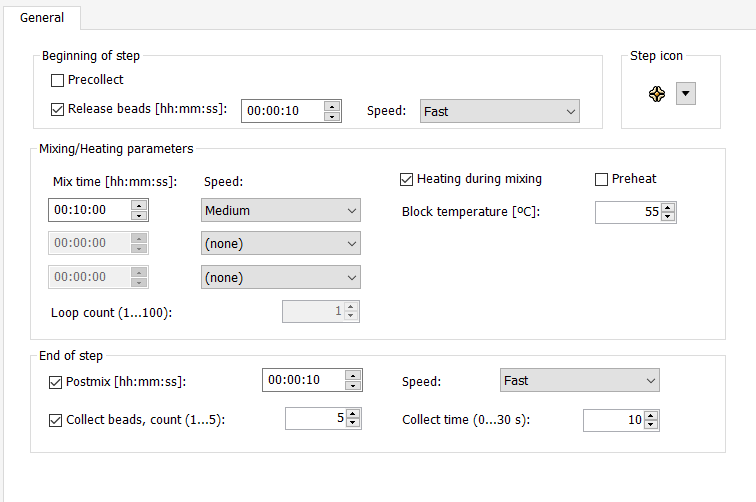

3) Add two washing steps to wash beads in washing buffer (steps ‘Wash 1’ and ‘Wash 2’; select plates ‘Wash 1’ and ‘Wash 2’ for each step, respectively. Parameters shown below are the same in both washing steps).

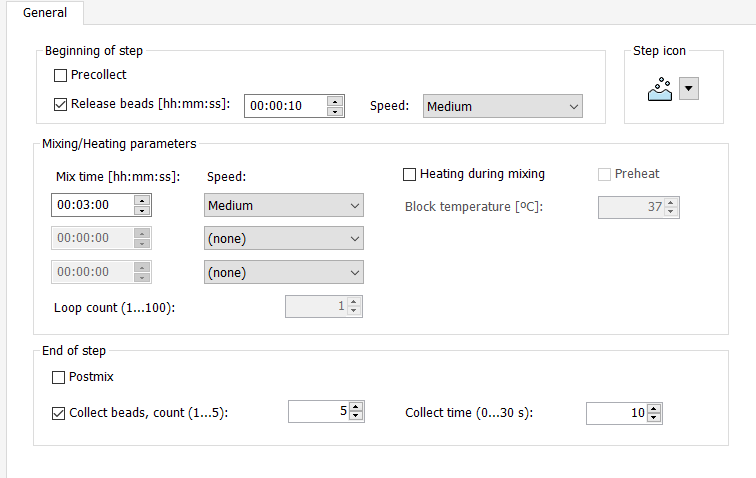

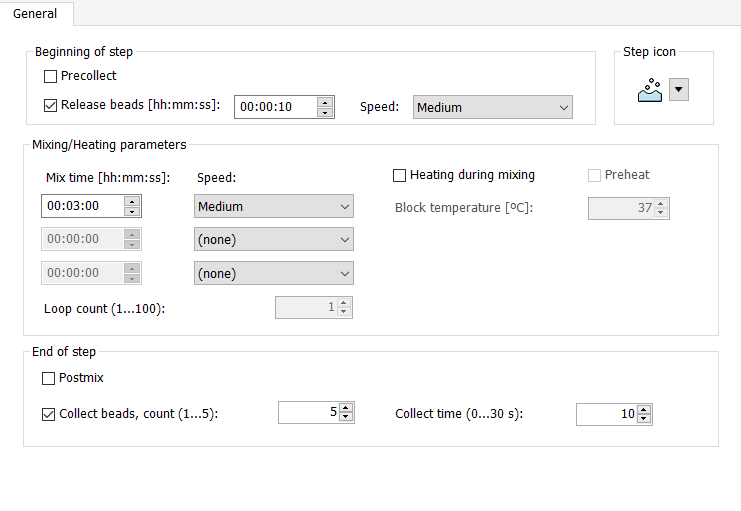
4) Add two washing steps to was beads in 80% ethanol (steps ‘Wash 3’ and ‘Wash 4’; select plates ‘Wash 3’ and ‘Wash 4’ for each step, respectively. Parameters below are the same for both washing steps).

5) Add the ‘Dry1’ step to airdry beads (select plate ‘Wash 4’).

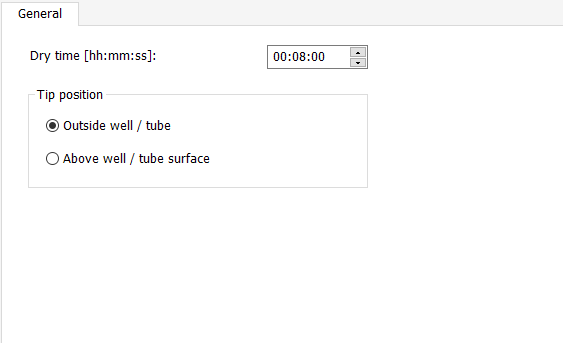

6) Add the ‘Elution’ step to elute DNA in C6 solution (select plate ‘Elution’).

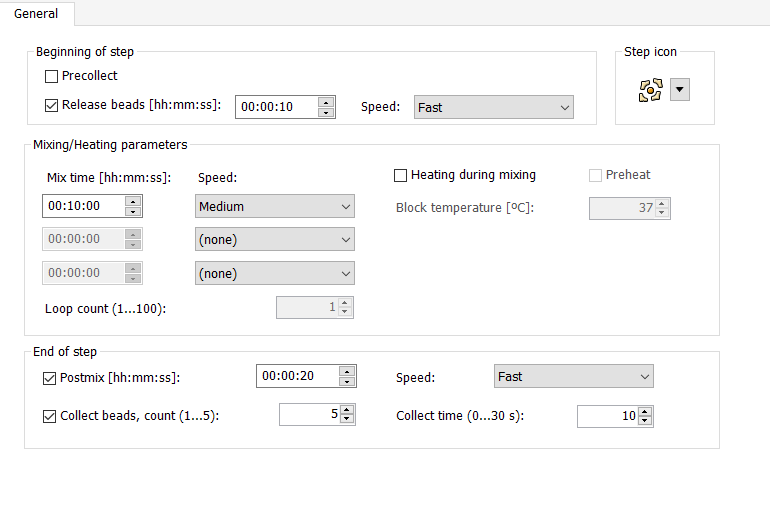

7) Add the ‘ReleaseBeads1’ step to release beads from tips (select plate ‘Wash 4’).

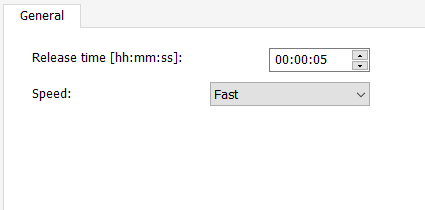

8) Leave tips in plate ‘Tip’.

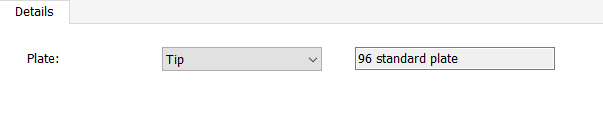
